# Co-infections by non-interacting pathogens are not independent & require new tests of interaction

**DOI:** 10.1101/618900

**Authors:** Frédéric M. Hamelin, Linda J.S. Allen, Vrushali A. Bokil, Louis J. Gross, Frank M. Hilker, Michael J. Jeger, Carrie A. Manore, Alison G. Power, Megan A. Rúa, Nik J. Cunniffe

## Abstract

If pathogen species, strains or clones do not interact, intuition suggests the proportion of co-infected hosts should be the product of the individual prevalences. Independence consequently underpins the wide range of methods for detecting pathogen interactions from cross-sectional survey data. However, the very simplest of epidemiological models challenge the underlying assumption of statistical independence. Even if pathogens do not interact, death of co-infected hosts causes net prevalences of individual pathogens to decrease simultaneously. The induced positive correlation between prevalences means the proportion of co-infected hosts is expected to be higher than multiplication would suggest. By modeling the dynamics of multiple non-interacting pathogens, we develop a pair of novel tests of interaction that properly account for non-independence. Our tests allow us to reinterpret data from previous studies including pathogens of humans, plants, and animals. Our work demonstrates how methods to identify interactions between pathogens can be updated using simple epidemic models.

## 1 Introduction

It is increasingly recognized that infections often involve multiple pathogen species or strains/clones of the same species (Balmer and Tanner, 2011; Vaumourin et al., 2015). Infection by one pathogen can affect susceptibility to subsequent infection by others (Griffiths et al., 2011; Petney and Andrews, 1998). Co-infection can also affect the severity and/or duration of infection, as well as the extent of symptoms and the level of infectiousness (Graham et al., 2005). Antagonistic, neutral and facilitative interactions are possible (Karvonen et al., 2018; Rigaud et al., 2010). Co-infection therefore potentially has significant epidemiological, clinical and evolutionary implications (Susi et al., 2015; Hilker et al., 2017; Alizon et al., 2013).

However, detecting and quantifying biological interactions between pathogens is notoriously challenging (Johnson and Buller, 2011; Hellard et al., 2015). In pathogens of some host taxa, most notably plant pathogens, biological interactions can be quantified by direct experimentation (Mascia and Gallitelli, 2016). However, often ethical considerations mean this is impossible, and so any signal of interaction must be extracted from population scale data. Analysis of longitudinal data remains the gold standard (Fenton et al., 2014), although the associated methods are not infallible (Telfer et al., 2010). However, collecting longitudinal data requires a dedicated and intensive sampling campaign, meaning in practice cross-sectional data are often all that are available. Methods for cross-sectional data typically concentrate on identifying deviation from statistical independence, using standard methods such as *χ*^2^ tests or log-linear modelling to test whether the observed probability of co-infection differs from the product of the prevalences of the individual pathogens (Booth and Bundy, 1995; Howard et al., 2001; Bogaert et al., 2004; Raso et al., 2004; Regev-Yochay et al., 2004; Nielsen et al., 2006; Chaturvedi et al., 2011; Rositch et al., 2012; Degarege et al., 2012; Malagón et al., 2016; Teweldemedhin et al., 2018). Detecting such a non-random statistical association between pathogens is then taken to signal a biological interaction. The underlying mechanism can range, for example, from individual-scale direct effects on within-host pathogen dynamics (Tollenaere et al., 2016; Mascia and Gallitelli, 2016), to indirect within-host immune-mediated interactions (de Roode et al., 2005), to indirect population-scale “ecological interference” caused by competition for susceptible hosts (Rohani et al., 1998, 2003).

A well-known difficulty is that factors other than biological interactions between pathogens can drive statistical associations. For instance, host heterogeneity – that some hosts are simply more likely than others to become infected – can generate positive statistical associations, since co-infection is more common in the most vulnerable hosts. Heterogeneity in host age can also generate statistical associations, as infections accumulate in older individuals (Lord et al., 1999; Kucharski and Gog, 2012; Kucharski et al., 2016). Methods aimed at disentangling such confounding factors have been developed, but show mixed results in detecting biological interactions (Pedersen and Fenton, 2007; Fenton et al., 2010; Hellard et al., 2012; Vaumourin et al., 2014). Methods using dynamic epidemiological models to track co-infections are also emerging, although more often than not require longitudinal data (Shrestha et al., 2011, 2013; Reich et al., 2013; Man et al., 2018; Alizon et al., 2019).

More fundamentally, however, the underpinning and long standing assumption that non-interaction implies statistical independence (Forbes, 1907; Cohen, 1973) has not been challenged. Here we confront the intuition that biological interactions can be detected via statistical associations, demonstrating how simple epidemiological models can change the way we think about biological interactions. In particular, we show that non-interacting pathogens should not be expected to have prevalences that are statistically independent. Co-infection by non-interacting pathogens is more probable than multiplication would suggest, invalidating any test invoking statistical independence.

The paper is organized as follows. First, we use a simple epidemiological model to show that the probability that a host is co-infected by both of a pair of non-interacting pathogens is greater than the product of the net prevalences of the individual pathogens. Second, we extend this result to an arbitrary number of non-interacting pathogens. This allows us to construct a novel test for biological interaction, based on testing the extent to which co-infection data can be explained by our epidemiological models in which pathogens do not interact. Different versions of this test, conditioned on the form of available data and whether or not co-infections are caused by different pathogen species, allow us to reinterpret a number of previous reports (Chaturvedi et al., 2011; López-Villavicencio et al., 2007; Andersson et al., 2013; Koepfli et al., 2011; Nickbakhsh et al., 2016; Seabloom et al., 2009; Moutailler et al., 2016; Howard et al., 2001; Molineaux et al., 1980). Our examples include plant, animal and human pathogens, and the methodology can potentially be applied to any cross-sectional survey data tracking co-infection.

## 2 Results

### 2.1 Two non-interacting pathogens

#### 2.1.1 Dynamics of the individual pathogens

We consider two distinct pathogen species, strains or clones (henceforth pathogens), which we assume do not interact, i.e. the interaction between the host and one of the pathogens is entirely unaffected by its infection status with respect to the other. Epidemiological properties that are therefore unaffected by the presence or absence of the other pathogen include initial susceptibility, within-host dynamics including rates of accumulation and/or movement within tissues, host responses to infection, as well as onward transmission. Assuming a fixed-size host population and Susceptible-Infected-Susceptible (S-I-S) dynamics (Keeling and Rohani, 2007), the proportion of the host population infected by pathogen *i* ∈ {1, 2} follows

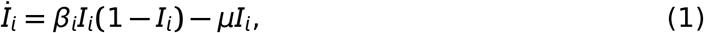

in which the dot denotes differentiation with respect to time, *β*_*i*_ is a pathogen-specific infection rate, and *μ* is the host’s natural death rate.

While natural mortality may be negligible for acute infections, it cannot be neglected for chronic (i.e. long-lasting) infections, which are responsible for a large fraction of co-infections in humans and animals (Griffiths et al., 2011; Gorsich et al., 2018). Likewise, plants remain infected over their entire lifetime following infection by most pathogens, including almost all plant viruses, as well as the anther smut fungus, which drives one of our examples here (López-Villavicencio et al., 2007).

We assume that the disease-induced death rate (virulence) is zero, as otherwise there would be ecological interactions between pathogens (Rohani et al., 2003). However our model can be extended to handle pathogen-specific rates of clearance (Supplementary Information: Sections S1.4, S1.5 and S2.3).

#### 2.1.2 Tracking co-infection

Making identical assumptions, but instead distinguishing hosts infected by different combinations of pathogens, leads to an alternate representation of the dynamics. We denote the proportion of hosts infected by only one of the two pathogens by *J*_*i*_, with *J*_1,2_ representing the proportion co-infected. Pathogen-specific net forces of infection are

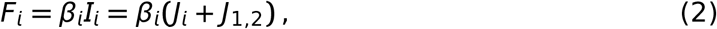

and so

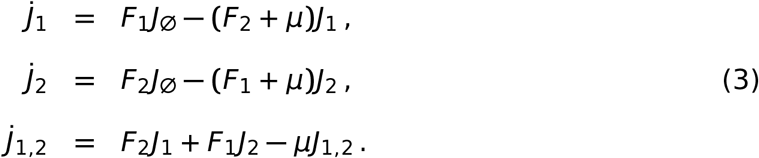

in which *J*_∅_ = 1 − *J*_1_ − *J*_2_ − *J*_1,2_ is the proportion of hosts uninfected by either pathogen (Fig. 1).

**Figure 1:**
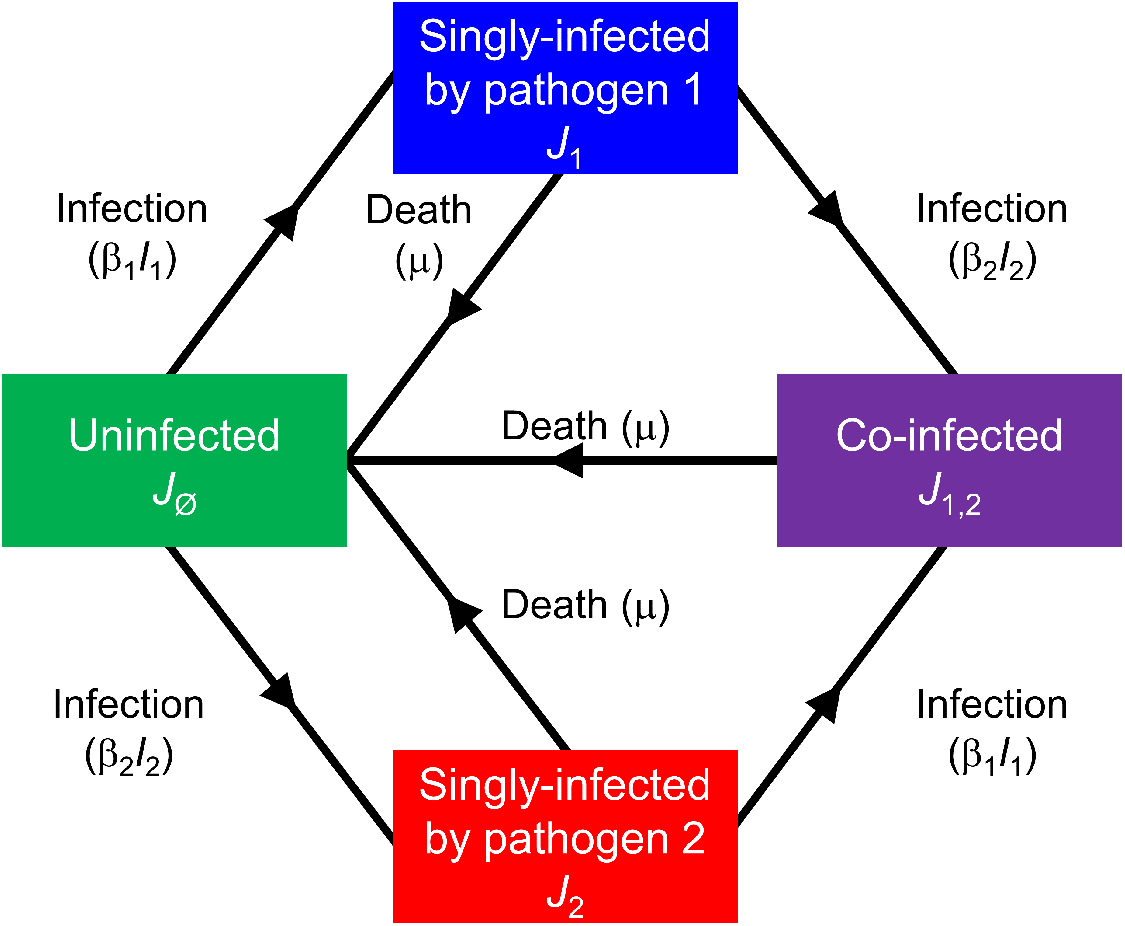
Schematic of the model tracking a pair of non-interacting pathogens. The model is defined in Eqs. (1–3): *J*_∅_ denotes uninfected hosts, *J*_1_ and *J*_2_ are hosts singly infected by pathogens 1 and 2, respectively, *J*_1,2_ are co-infected hosts, *I*_1_ =*J*_1_ +*J*_1,2_ and *I*_2_ =*J*_2_ +*J*_1,2_ are net densities of hosts infected by pathogens 1 and 2, respectively.

#### 2.1.3 Prevalence of co-infected hosts

We assume the basic reproduction number, *R*_0,*i*_ = *β*_i_/*μ* > 1 for both pathogens. Solving Eq. (3) numerically for arbitrary but representative parameters (Fig. 2A) shows the proportion of co-infected hosts (*J*_1,2_) to be larger than the product of the individual prevalences (*P* = *I*_1_*I*_2_ from Eq. (1)). That *J*_1,2_(*t*) ≥ *P*(*t*) for large *t* (for all parameters) can be proved analytically (Supplementary Information: Section S1.1).

**Figure 2:**
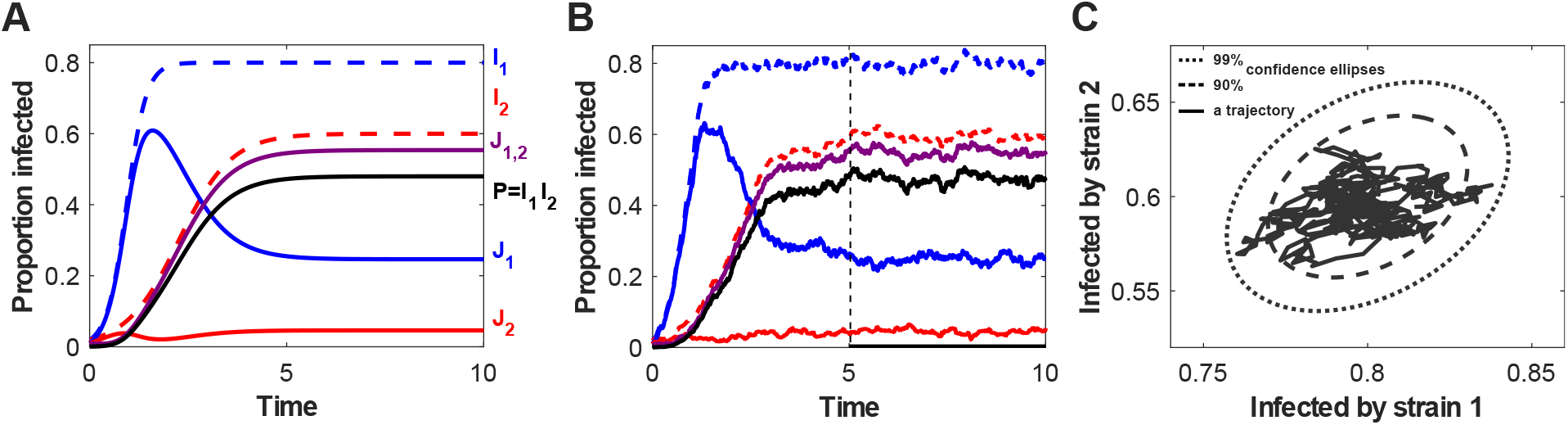
Simulations of the two-pathogen model show that net densities of the two pathogens are positively correlated. *J*_1_ and *J*_2_ are hosts singly infected by pathogens 1 and 2, respectively, *J*_1,2_ are co-infected hosts, *I*_1_ = *J*_1_ + *J*_1,2_ and *I*_2_ = *J*_2_ + *J*_1,2_ are net densities of hosts infected by pathogens 1 and 2, respectively. (A) Dynamics of the deterministic model (1-3), with *β*_1_ = 5, *β*_2_ = 2.5, and *μ* = 1 (parameters have units of inverse time). (B) Dynamics of a stochastic version of the model, in a population of size *N* = 1000 (see also Methods: Section 4.1.4, “Stochastic models”). (C) A single trajectory from the stochastic simulation (black line) in panel B (restricted to the time interval starting from the dashed line at *t* = 5) in the phase plane (*I*_1_, *I*_2_ ), and the 90% and 99% confidence ellipses (dashed and dotted curves, respectively) generated from an analytical approximation to the stochastic model.

Simulations of a stochastic analogue of the model (Fig. 2B) reveal the key driver of this behavior. The net prevalences of the pathogens considered in isolation, *I*_1_ and *I*_2_, are positively correlated (Fig. 2C; Eq. (27) in Methods: Section 4.1.4, “Stochastic models”), due to simultaneous reductions whenever co-infected hosts die.

#### 2.1.4 Deviation from statistical independence

For *R*_0,*i*_ > 1 the relative deviation of the equilibrium prevalence of co-infection 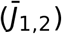 from that required by statistical independence 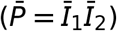 is

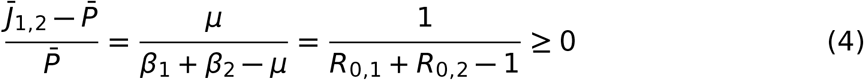

(Eq. (9) in Methods: Section 4.1.1, “Equilibria of the two-pathogen model”). The deviation is therefore zero if and only if the host natural death rate *μ* = 0.

The observed outcome would conform with statistical independence only for non-interacting pathogens where there is no host natural death (at the timescale of an infection). This reiterates the role of host natural death in causing deviation from a statistical association pattern.

This result (Eq. (4)) was first published by Kucharski and Gog (2012) in a different context (model reduction in multi-strain influenza models). Moreover, using a continuous age-structured model, Kucharski and Gog (2012) showed that one may recover statistical independence within infinitesimal age-classes. The result in Eq. (4) is related to aging, as individuals acquire more infections as they age. As age increases, so does the probability of being infected with pathogens 1 and/or 2. Therefore, the prevalences of pathogens 1 and 2 are positively correlated (Kucharski et al., 2016). The reason for a greater deviation from independence as the mortality rate *μ* increases is likely due to the fact that prevalence is increasing and concave with respect to age, and saturates in older age-classes (Lord et al., 1999).

### 2.2 Testing for interactions between pathogens

Eq. (3) can be straightforwardly extended to track *n* pathogens which do not interact in any way (including pairwise and three-way interactions). Equilibria of this model are prevalences of different classes of infected or co-infected hosts carrying different combinations of non-interacting pathogens. These can be used to derive a test for interaction between pathogens which properly accounts for the lack of statistical independence revealed by our analysis of the simple two-pathogen model.

#### 2.2.1 Modelling co-infection by *n* non-interacting pathogens

We denote the proportion of hosts simultaneously co-infected by the (non-empty) set of pathogens Γ to be *J*_Γ_, and use Ω_*i*_ = Γ \ {*i*} (for *i* ∈ Γ) to represent combinations with one fewer pathogen.

The dynamics of the 2^*n*^ − 1 distinct values of *J*_Γ_ follow

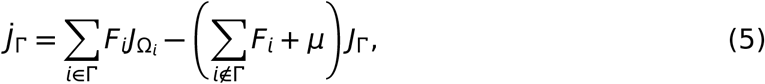

in which the net force of infection of pathogen *i* is

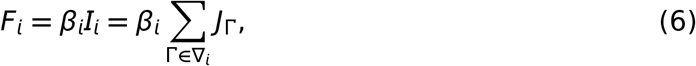

and ∇_*i*_ is the set of all subsets of {1, …, *n*} containing *i* as an element. Eq. (5) can be interpreted by noting 
- the first term tracks inflow due to hosts carrying one fewer pathogen becoming infected;
- the second term tracks the outflows due to hosts becoming infected by an additional pathogen, or death.

##### Predicted prevalences

If *R*_0,*i*_ = *β*_*i*_/*μ* > 1 for all *i* = 1, …, *n*, the equilibrium prevalence of hosts infected by any given combination of pathogens, 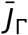, can be obtained by (recursively) solving a system of 2^*n*^ linear equations (Eq. (16) in Methods: Section 4.1.2, “Equilibria of the *n*-pathogen model”).

These equilibrium prevalences are the prediction of our “Non-interacting Distinct Pathogens” (NiDP) model, which in dimensionless form has *n* parameters (the *R*_0,*i*_’s, *i* = 1, …, *n*; Methods: Section 4.2.2, “Fitting the models”).

##### Epidemiologically interchangeable pathogens

If we simplify the model by assuming that all pathogen infection rates are equal (i.e. *β*_*i*_ = *β* for all *i*), then if *R*_0_ = *β*/*μ* > 1, the proportion of hosts infected by *k* distinct pathogens can be obtained by (recursively) solving *n* +1 linear equations (Eq. (22) in Methods: Section 4.1.3, “Deriving the NiSP model from the NiDP model”). This constitutes the prediction of our “Non-interacting Similar Pathogens” (NiSP) model, a simplified form of the NiDP model requiring only a single parameter (*R*_0_).

#### 2.2.2 Using the models to test for interactions

If either the NiSP or NiDP model adequately explain co-infection data, those data are consistent with the underpinning assumption that pathogens do not interact. Which model is fitted depends on the form of the available data.

##### Numbers of distinct pathogens

Studies often quantify only the number of distinct pathogens carried by individual hosts, without necessarily specifying the combinations involved (López-Villavicencio et al., 2007; Seabloom et al., 2009; Chaturvedi et al., 2011; Koepfli et al., 2011; Andersson et al., 2013; Moutailler et al., 2016; Nickbakhsh et al., 2016). There are insufficient degrees of freedom in such data to fit the NiDP model, and so we fall back upon the NiSP model. In using the NiSP model, we additionally assume all pathogens within a given study are epidemiologically inter-changeable.

We identified four suitable studies reporting data concerning strains/clones of a single pathogen, and tested whether these data are consistent with no interaction. For all four studies (Fig. 3), the best-fitting NiSP model is a better fit to the data than the corresponding binomial model assuming statistical independence (Eq. (28) in Methods: Section 4.2.1, “Models corresponding to assuming statistical independence”). Three additional examples for data-sets considering distinct pathogens, which deviate more markedly from the epidemiological equivalence assumption, are in the Supplementary Information (Section S2.1).

**Figure 3:**
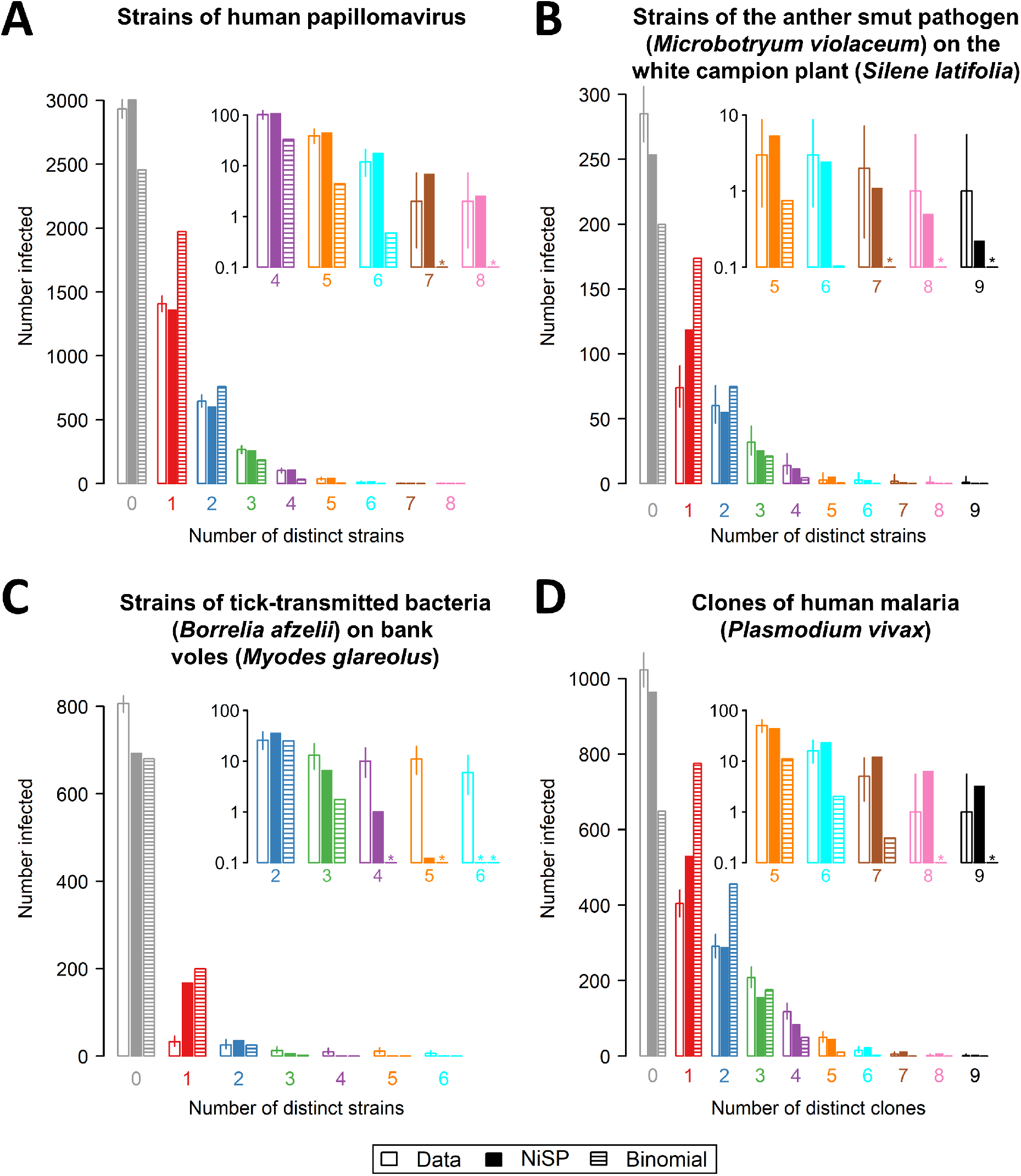
Comparing predictions of the NiSP model with binomial models assuming statistical independence. In using the NiSP model, pathogens are assumed to be epidemiologically interchangeable: we have therefore restricted attention to data sets concerning strains/clones of a single pathogen species. (A) strains of human papillomavirus (Chaturvedi et al., 2011); (B) strains of the anther smut pathogen (*M. violaceum*) on the white campion (*S. latifolia*) (López-Villavicencio et al., 2007); (C) strains of tick-transmitted bacteria (*B. afzelii*) on bank voles (*M. glareolus*) (Andersson et al., 2013); and (D) clones of malaria (*P. vivax*) (Koepfli et al., 2011). Insets to each panel show a “zoomed-in” section of the graph corresponding to high multiplicities of clone/strain co-infection. Asterisks indicate predicted counts smaller than 0.1. In all four cases, the NiSP model is a better fit to the data than the binomial model (ΔAIC = 572.8, 158.6, 293.8 and 596.3, respectively). For the data shown in panel (A), there is no evidence that the NiSP model does not fit the data (lack of goodness-of-fit *p* = 0.08), and so our test indicates the human papillomavirus strains do not interact. For the data shown in panels (B)-(D), there is evidence of lack of goodness-of-fit (all have lack of goodness-of-fit *p* < 0.01). Our test therefore indicates these strains/clones interact (or are epidemiologically different).

In one case – co-infection by different strains of human papillomavirus (Chaturvedi et al., 2011) (Fig. 3A) – we find no evidence that the reported data cannot be explained by the NiSP model. These data therefore support the hypothesis of no inter-action – and indeed no epidemiological differences – between the pathogen strains in question.

In the three other cases we considered – strains of anther smut (*Microbotryum violaceum*) on the white campion (*Silene latifolia*) (López-Villavicencio et al., 2007) (Fig. 3B); strains of the tick-transmitted bacterium *Borrelia afzelii* on bank voles (*Myodes glareolus*) (Andersson et al., 2013) (Fig. 3C); and clones of a single malaria parasite (*Plasmodium vivax*) infecting children (Koepfli et al., 2011) (Fig. 3D) – despite outperforming the model corresponding to statistical independence, the best-fitting NiSP model does not adequately explain the data. We therefore reject the hypotheses of no interaction in all three cases, noting that our use of the NiSP model means it might be epidemiological differences between pathogen strains/clones that have in fact been revealed.

##### Combinations of pathogens

Other studies report the proportion of hosts infected by particular combinations (rather than counts) of pathogens, although many of those concentrate on helminth macroparasites for which our underlying S-I-S model is well-known to be inappropriate (Anderson and May, 1991).

However, a methodological article by Howard et al. (2001) introduces the use of log-linear modeling to test for statistical associations. Conveniently, that article reports the results of that methodology as applied to a large number of studies focusing on *Plasmodium* spp. causing malaria.

By interrogating the original data sources (Methods: Section 4.3.2, “Combinations of pathogens (NiDP model)”), we found a total of 41 studies of malaria reporting the disease status of at least *N* = 100 individuals, and in which three of *P. falciparum*, *P. malariae*, *P. ovale* and *P. vivax* were considered. Data therefore consist of counts of the number of individuals infected with different combinations of three of these four pathogens, a total of eight classes. There were sufficient degrees of freedom to fit the NiDP model, which here has three parameters, each corresponding to the infection rate of a single *Plasmodium* spp. Fig. 4A shows the example of fitting the NiDP model to data from a study of malaria in Nigeria (Molineaux et al., 1980).

**Figure 4:**
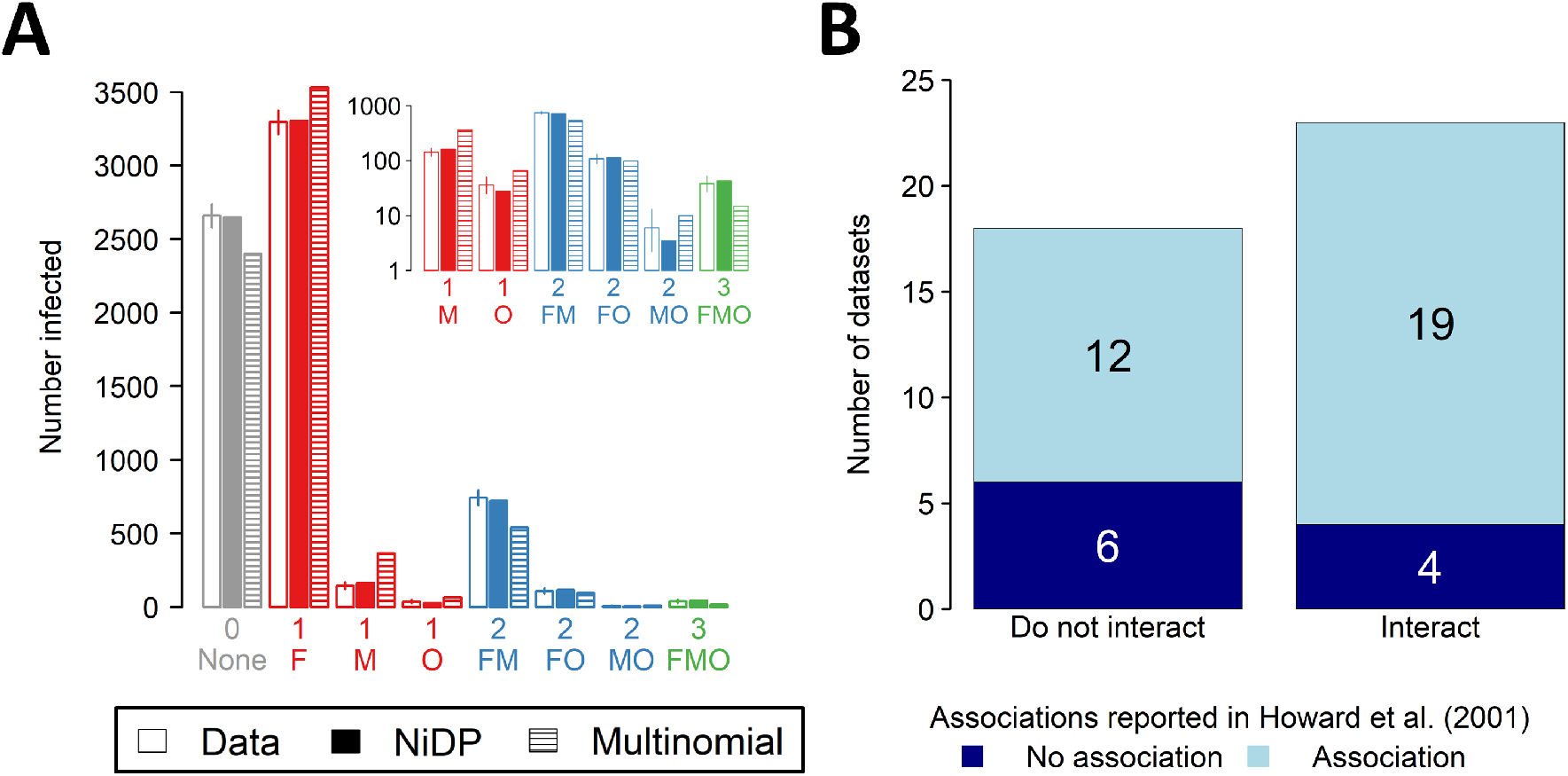
Using the NiDP model to re-analyse malaria data sets considered by Howard et al. (2001). In using the NiDP model there is no need to assume malaria-causing *Plasmodium* spp. are epidemiologically interchangeable. (A) Comparing the predictions of the NiDP model with a multinomial model of infection (i.e. statistical independence) for the data set on *P. falciparum* (F), *P. malariae* (M) and *P. ovale* (O) co-infection in Nigeria reported by Molineaux et al. (1980). The NiDP model is a better fit to the data than the multinomial model (ΔAIC = 326.2); additionally, there is no evidence of lack of goodness-of-fit (*p* = 0.40). This data set is therefore consistent with no interaction between the three *Plasmodium* species. (B) Comparing the results of fitting the NiDP model and the methodology of Howard et al. (2001) based on log-linear regression and so statistical-independence. For 16 (i.e. 12 + 4) out of the 41 data sets we considered, the conclusions of the two methods differ.

Fitting the NiDP model allows us to test for interactions between *Plasmodium* spp., without assuming they are epidemiologically interchangeable. In 18 of the 41 cases we considered, our methods suggest the data are consistent with no interaction (Fig. 4B). We note that in 12 of these 18 cases the methodology based on statistical independence of Howard et al. (2001) instead suggests the *Plasmodium* spp. interact.

## 3 Discussion

We have shown that pathogens which do not interact and so have uncoupled prevalence dynamics (Eq. (1)) are not statistically independent. For two pathogens, the prevalence of co-infection is always greater than the product of the prevalences (Eq. (4)), unless host natural death does not occur. This result was first published in an age-structured, multi-strain influenza model (Kucharski and Gog, 2012). Pathogens share a single host in co-infections, and so when a co-infected host dies, net prevalences of both pathogens decrease simultaneously. The prevalences of individual pathogens, regarded as random variables, therefore co-vary positively. A related interpretation is due to Kucharski and Gog (2012): the prevalences of the pathogens are positively correlated through a single independent variable, namely the age of the hosts. As a side result, we note our analysis indicates a high-profile model of May and Nowak (1995) is based on a faulty assumption of probabilistic independence (Supplementary Information: Section S1.3). More importantly, our analysis shows that statistically independent pathogens may well be interacting (Supplementary Information: Section S1.5) which confirms that statistical independence is far from equivalent to the absence of biological interaction between pathogens.

We extended our model to an arbitrary number of pathogens to develop a novel test for interaction that properly accounts for statistical non-independence. Many data sets summarize co-infections in terms of multiplicity of infection, regardless of which pathogens are involved. Since there would be as many epidemiological parameters as pathogens in our default NiDP model, and so as many parameters as data-points, the full model would be over-parameterised. We therefore introduced the additional assumption that all pathogens are epidemiologically interchangeable. This formed the basis of the parsimonious NiSP model, which is most appropriate for testing for interactions between strains or clones of a single pathogen species.

Despite the strong and perhaps even unrealistic assumption that strains/clones are interchangeable, the NiSP model outperformed the binomial model assuming statistical independence for all four data sets we considered. In particular, the NiSP model successfully captured the fat tails characteristic of observed multiplicity of infection distributions. All four data sets therefore support the idea that co-infection is far more frequent than statistical independence would imply.

For the data set concerning co-infection by different strains of human papillomavirus (Chaturvedi et al., 2011), the NiSP model also passed the goodness-of-fit test, allowing us to conclude strains of this pathogen do not interact. Goodness-of-fit for such a simple model is a particularly conservative test, especially for the NiSP model, when we assume pathogens clones/strains are epidemiologically interchangeable.

We illustrated our methods via case studies for which suitable data are readily available, and our purpose was not to come to definitive conclusions concerning any particular system. That would require dedicated studies. However, by fitting even a highly-simplified version of our model to data, we have demonstrated how results of simple epidemiological models challenge previous methods based on statistical independence.

To explore further the implications of our findings, we analyzed available data sets tracking combinations of pathogens involved in each occurrence of co-infection. For methodological comparison purposes, we restricted ourselves to data referenced in Howard et al. (2001), concerning interactions between *Plasmodium* spp. causing malaria. Relaxing the assumption of epidemiological interchangeability (i.e. using the NiDP model), we found that 43.9% (i.e. 18/41) of data sets considered in Howard et al. (2001) are consistent with no interaction.

One may wonder whether focusing on age classes may be sufficient to correct for the positive correlation between non-interacting pathogens (Lord et al., 1999). Of the 41 data sets identified by Howard et al. (2001) that we analysed, 14 focused only on data collected from children, and therefore, associations are less likely less likely to emerge solely by the confounding effect of age (Fenton et al., 2010). Of these 14 studies, we came to the same conclusion as Howard et al. (2001) in only 6 cases. We identified 2 cases in which our methods suggest there is an interaction in which Howard et al. (2001) concluded no interaction (studies 71 and 77), as well as 6 cases in which we conclude no interaction whereas Howard et al. (2001) conclude there is an interaction (studies 76, 68, 69, 70, 79 and 80). Thus, focusing on discrete and arbitrary age classes may not be sufficient to correct for the positive correlation between non-interacting pathogens.

Again we do not intend to conclusively demonstrate interactions – or lack of interactions – for malaria. Instead what is important is that our results very often diverge from those originally reported in Howard et al. (2001) using a method based on statistical associations, namely log-linear regression. Log-linear regression suffers from well-acknowledged difficulties in cases in which there are zero counts (i.e. certain combinations of pathogens are not observed) (Fienberg and Rinaldo, 2012). Such cases often arise in epidemiology. Methods based on epidemiological models therefore offer a twofold advantage: biological interactions are not confounded with statistical associations, and parameter estimation is well-posed, irrespective of zero counts.

Moreover, simple epidemiological models (with no explicit age structure) intrinsically correct the bias due to the positive correlation between age and prevalence, which makes it unnecessary to control for age. Therefore, and this may be our main conclusion: although age is an evident confounding factor, epidemiological models make it unnecessary to keep track of the age of infected hosts. This is made possible by replacing the paradigm of “statistical Independence and random distributions” with “model-based distributions in absence of biological interactions.”

We focused here on the simple S-I-S model, since it is sufficiently generic to be applicable to a number of systems. However, an important assumption of our model is that natural mortality occurs at a time scale comparable to that of an infection. Our model is therefore tailored for chronic (i.e. long-lasting) infections, which represent a large fraction of of co-infections in humans, animals, and plants. Also, our study is restricted to nonlethal infections, as otherwise there may be ecological interactions between pathogens. In future work focusing on particular pathogens, models including additional system-specific detail would of course be appropriate. We leave further analysis of more complex underpinning epidemiological models to such future research.

Lastly, we speculate our results may have implications beyond epidemiology. After all, pathogens are species which form meta-populations occupying discrete patches (hosts) (Seabloom et al., 2015). Meta-community ecology has long been concerned with whether interactions between species can be detected from co-occurrence data (Forbes, 1907; Caswell, 1976; Connor and Simberloff, 1979) and most existing methods are based on detecting statistical associations (Gotelli, 2000; Gotelli and Ulrich, 2012), but see Hastings (1987). Our dynamical modeling approach may also provide a new perspective in this area.

## 4 Methods

### 4.1 Mathematical analyses

#### 4.1.1 Equilibria of the two-pathogen model

The 2-pathogen model is given by Eq. (1–2–3). Since the population size is constant, *J*_∅_ = 1 − *J*_1_ − *J*_2_ − *J*_1,2_, and so it follows that

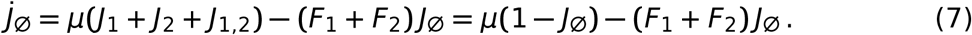

It is well-known (Keeling and Rohani, 2007) that if *R*_0,*i*_ = *β*_*i*_/*μ* > 1 and *I*_*i*_(0) > 0, the prevalence of pathogen *i* will tend to an equilibrium 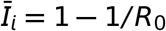.

Since *F*_*i*_ = *β*_*i*_*I*_*i*_ and *J*_*i*_ = *I*_*i*_ − *J*_1,2_, the rate of change of co-infected hosts in Eq. (3) can be recast as

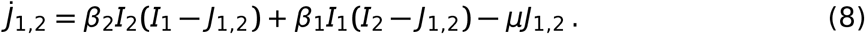

The equilibrium prevalence of co-infected hosts 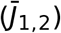 can therefore be written in terms of the individual net prevalences at equilibrium (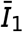 and 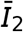),

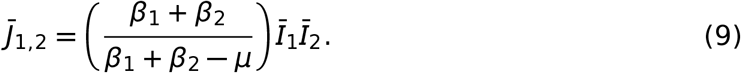

This immediately leads to the result concerning the deviation of 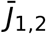 from 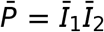 (i.e., the expected prevalence of co-infected hosts given statistical independence) quoted in Eq. (4).

#### 4.1.2 Equilibria of the *n*-pathogen model

The *n*-pathogen model is given by Eq. (1–5–6). Since the host population size is constant, *J*_∅_ = 1 − Σ_Γ∈∇_*J*_Γ_, where ∇ is the set of all 2^*n*^ − 1 sets with infected or co-infected hosts. It is also true that

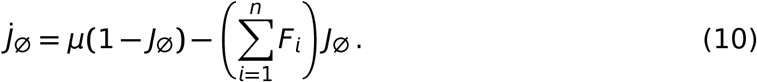

At equilibrium, Eq. (5) becomes

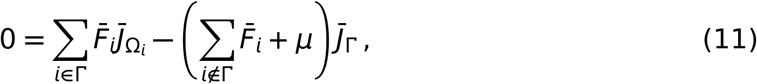

in which 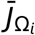 and 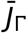 are equilibrium prevalences, and 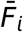 is the force of infection of pathogen *i* at equilibrium, i.e.

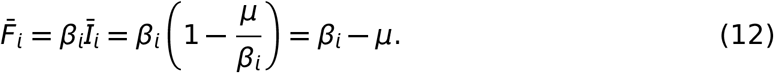

Since these forces of infection are constant and do not depend on the equilibrium prevalences, the set of 2^*n*^ − 1 equations partially characterizing the equilibrium is linear, with

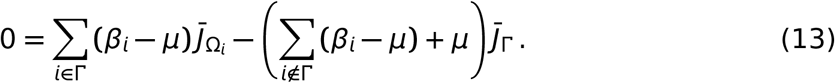

Similarly, Eq. (10) is linear

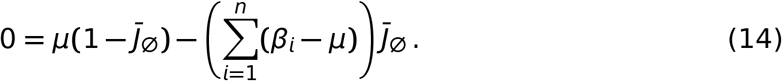

The equilibrium prevalences can be written very conveniently in a recursive form (i.e. using the first equation to fix 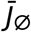, using 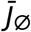 to independently calculate all values of 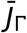 for |Γ| = 1, then using the set of values of 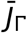 when |Γ| = 1 to independently calculate all values of 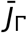 for |Γ| = 2, and so on). A recurrence relation to find all equilibrium prevalences can therefore be initiated with the following expression for the density of uninfected hosts:

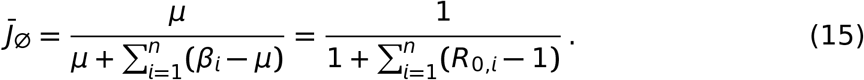

Then, one may recursively use the following equation, equivalent to Eq. (13):

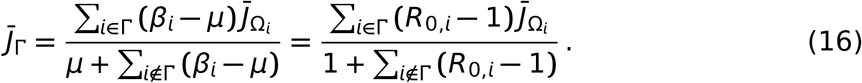

Since the densities in Eq. (16) are entirely in terms of the equilibrium densities of hosts carrying one fewer pathogen 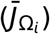, this allows us to recursively find the densities of all pathogens given pathogen-by-pathogen values of *R*_0,*i*_.

#### 4.1.3 Deriving the NiSP model from the NiDP model

If all pathogens are interchangeable, and so have identical values of *R*_0,*i*_ = *R*_0_ ∀*i*, then for any pair of combinations of infecting pathogens, Γ_1_ and Γ_2_, it must be the case that 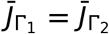 whenever |Γ_1_| = |Γ_2_|. This means the equilibrium prevalences of hosts infected by the same number of distinct pathogens must all be equal, irrespective of the particular combination of pathogens that is carried. In this case solving the system is much simpler. First, Eq. (11) can be rewritten as

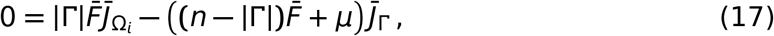

in which 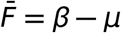. The net prevalence of hosts infected by *k* distinct pathogens is

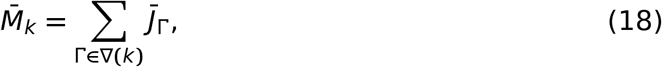

in which ∇(*k*) is the set of combinations of {1, …, *n*} with *k* elements. Since the form of Eq. (17) depends only on |Γ|, all individual prevalences involved in 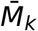 are identical, and so

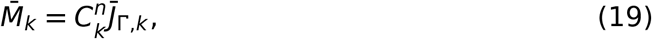

in which 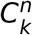 is a combinatorial coefficient, and 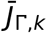 is any of the individual prevalences for which |Γ| = *k*. The ratio between successive values of 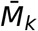 is given by

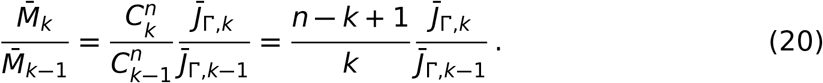

From Eq. (15), it follows that

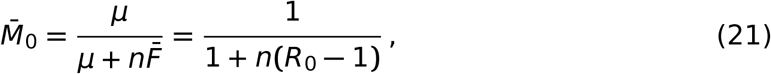

in which *R*_0_ = *β*/*μ*. For 1 ≤ *k* ≤ *n*, Eq. (17) and (20) together imply

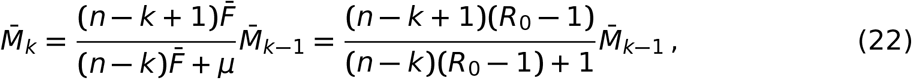

a form which admits a simple recursive solution.

#### 4.1.4 Stochastic models

Figs. 2B and 2C were generated by simulating the stochastic differential equation corresponding to Eq. (3); simulating a continuous time Markov chain model using Gillespie’s algorithm gave consistent results. Confidence ellipses were obtained from an approximate expression for the covariance matrix at equilibrium (see below).

##### Continuous-time Markov chain

The continuous-time Markov chain model corresponding to the unscaled version of Eq. (3–7) tracks a vector of integer-valued random variables *X*(*t***) = (***J*_∅_(*t*), *J*_1_(*t*), *J*_2_(*t*), *J*_1,2_(*t*)). Defining Δ*X* = *X*(*t* + Δ*t*) − *X*(*t***) = (**Δ*J*_∅_, Δ*J*_1_, Δ*J*_2_, Δ*J*_1,2_), changes of ±1 to each element of *X*(*t*) occur in small periods of time Δ*t* at the rates given in Table 1. Stochastic trajectories from this model can conveniently be simulated via the Gillespie algorithm (Gillespie, 1977). Note that the numeric values of the infection rates and the host birth rate must be altered to account for the scaling by population size.

**Table 1:** Transitions in the two-pathogen stochastic models. The prevalence of uninfected host is *J*_∅_, the prevalence of each class of singly-infected hosts is *J*_*i*_ (for *i* ∈ [1, 2]), and the prevalence of co-infected host is *J*_1,2_. The net force of infection of pathogen *i* is *F*_*i*_ = *β*_*i*_*I*_*i*_/*N* = *β*_*i*_(*J*_*i*_ + *J*_1,2_ )/*N* (note the scaling by the population size *N* relative to the forces of infection as used in the deterministic version of the model). To ensure a constant host population size, we have made the simplifying assumption that removal and replacement occur simultaneously; this has no effect on our qualitative results.

| Event number | Event | Rate | Change(s) to state variable(s) ($\Delta X$) |
| --- | --- | --- | --- |
| 1 | Infection of uninfected host by pathogen 1 | $F_1 J_\emptyset \Delta t + o(\Delta t)$ | $J_\emptyset \rightarrow J_\emptyset - 1$<br>$J_1 \rightarrow J_1 + 1$ |
| 2 | Infection of uninfected host by pathogen 2 | $F_2 J_\emptyset \Delta t + o(\Delta t)$ | $J_\emptyset \rightarrow J_\emptyset - 1$<br>$J_2 \rightarrow J_2 + 1$ |
| 3 | Infection by pathogen 1 of host singly-infected by pathogen 2 | $F_1 J_2 \Delta t + o(\Delta t)$ | $J_2 \rightarrow J_2 - 1$<br>$J_{1,2} \rightarrow J_{1,2} + 1$ |
| 4 | Infection by pathogen 2 of host singly-infected by pathogen 1 | $F_2 J_1 \Delta t + o(\Delta t)$ | $J_1 \rightarrow J_1 - 1$<br>$J_{1,2} \rightarrow J_{1,2} + 1$ |
| 5 | Death of host singly-infected by pathogen 1 and replacement with an uninfected host | $\mu J_1 \Delta t + o(\Delta t)$ | $J_1 \rightarrow J_1 - 1$<br>$J_\emptyset \rightarrow J_\emptyset + 1$ |
| 6 | Death of host singly-infected by pathogen 2 and replacement with an uninfected host | $\mu J_2 \Delta t + o(\Delta t)$ | $J_2 \rightarrow J_2 - 1$<br>$J_\emptyset \rightarrow J_\emptyset + 1$ |
| 7 | Death of co-infected host and replacement with an uninfected host | $\mu J_{1,2} \Delta t + o(\Delta t)$ | $J_{1,2} \rightarrow J_{1,2} - 1$<br>$J_\emptyset \rightarrow J_\emptyset + 1$ |

##### Stochastic differential equations

The model can also be written as a system of stochastic differential equations (SDEs), an approximation to the continuous-time Markov chain that is valid for sufficiently large *N* (Kurtz, 1970) and which is particularly well-suited for simulation of the stochastic model when the population size is large. This form of the model again tracks the seven events in Table 1, although in the SDE formulation the random variables in *X*(*t*) are continuous-valued. A heuristic derivation is based on a normal approximation described below. Alternately, the forward Kolmogorov differential equations in the continuous-time Markov chain model are closely related to the Fokker Planck equation for the probability density function of the SDE model (Allen et al., 2008).

The expected change 𝔼(Δ*X*) and covariance of the changes 𝕍(Δ*X*) can be computed from Table 1 to order Δ*t* via

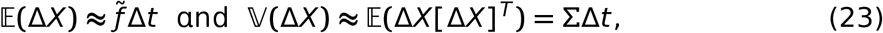

where 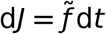 is the unscaled version of the deterministic model as specified in Eq. (3–7) with *N* = *J*_∅_ + *J*_1_ + *J*_2_ +*J*_1,2_ (a constant) and *F*_*i*_ = *β*_*i*_(*J*_*i*_ + *J*_1,2_)/*N*. In addition, the matrix Σ is given by

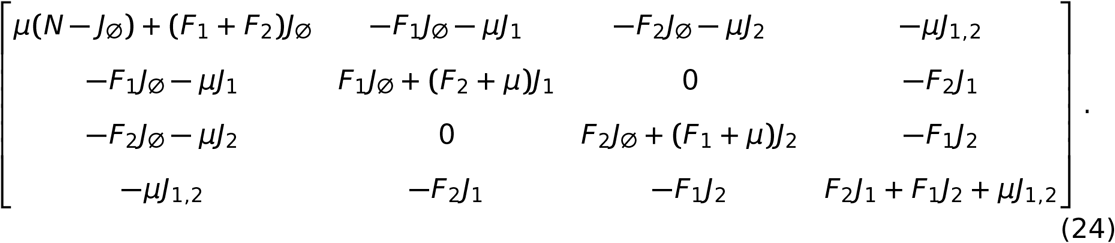

The changes in a small time interval Δ*t* are approximated by a normal distribution via the Central Limit Theorem: Δ*X*(*t*) − 𝔼(Δ*X*(*t*)) ≈ Normal(0, ΣΔ*t*), where 0 = zero vector. The covariance matrix Σ can be written as Σ = *GG*^*T*^. Letting Δ*t* → 0, the SDE model can therefore be expressed as

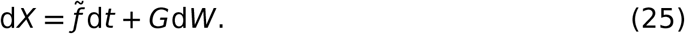

The matrix *G* is not unique but a simple form with dimension 4 × 7 accounts for each event in Table 1 (Allen et al., 2008). Each entry in matrix *G* involves a square root and *W* is a vector of seven independent standard Wiener processes, where d*W*_*i*_ ≈ Δ*W*_*i*_(*t*) = *W*_*i*_(*t* + Δ*t*) − *W*_*i*_(*t*) ~ Normal(0, Δ*t*). An explicit form for the SDE model in Eq. (25) is

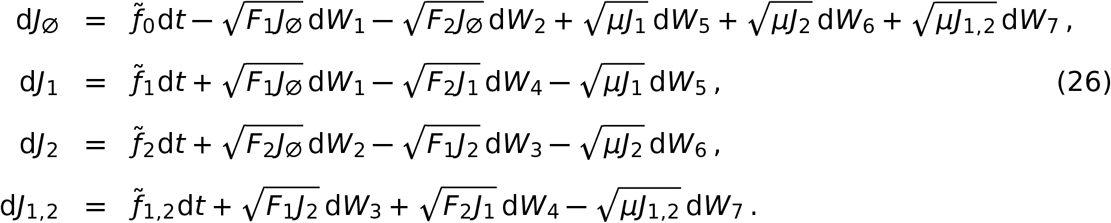

##### Covariance matrix at the endemic equilibrium

In Supplementary Information (Section S1.2) we show that the covariance between the prevalences of pathogen 1 and pathogen 2 as they fluctuate in the vicinity of their equilibrium values is approximately

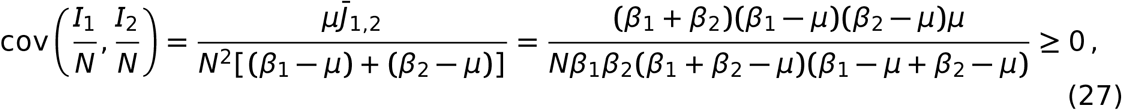

with equality if and only if *μ* = 0 (assuming *β*_*i*_ > *μ*, *i* = 1, 2). Only in the specific case *μ* = 0, is the deviation from statistical independence equal to zero (Eq. (4)).

### 4.2 Statistical methods

#### 4.2.1 Models corresponding to assuming statistical independence

If data are observations of numbers of individuals infected with *k* distinct pathogens, *O*_*k*_, for *k* ∈ [0, *n*], statistical independence corresponds to assuming the infection load of a single individual follows the one-parameter, binomial model *Bin*(*n*, *p*), in which *p* is the pathogen prevalence (assumed identical for each pathogen, and fitted appropriately to the data), and *n* is the maximum number of infections that is possible (i.e. the total number of distinct pathogens under consideration). Model predictions are then simply *N* samples from this binomial distribution, where *N* = Σ_*k*_ *O*_*k*_ is the total number of individuals observed in the data. One interpretation is as a multinomial model in which

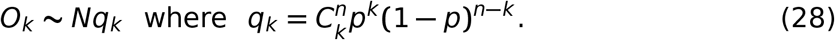

For the data for malaria corresponding to numbers of individuals, *O*_Γ_, infected by different sets of pathogens, Γ, statistical independence corresponds to an *n*-parameter multinomial model, parameterised by the prevalences of the individual pathogens *p_i_* (again fitted to the data), i.e.

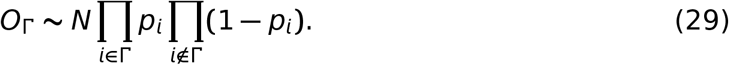

#### 4.2.2 Fitting the models

The host natural death rate, *μ*, can be scaled out of the equilibrium prevalences by rescaling time. Fitting the models therefore corresponds to finding value(s) for scaled infection rate(s) *β*_*i*_, i.e. *R*_0,*i*_ = *β*_*i*_/*μ* (all are equal for the NiSP model).

The method used to fit the model does not depend on whether the data are numbers of hosts infected by a particular combination of pathogens, or numbers of hosts carrying particular numbers of distinct pathogens, since both can be viewed as *N* samples drawn from a multinomial distribution, with *q*_*j*_ observations of the *j*^*th*^ class. If the corresponding probabilities generated by the model being fitted are *p*_*j*_, then the log-likelihood is

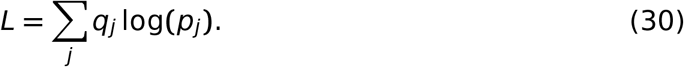

The models were fitted by maximizing *L* via optim() in R (R Core Team, 2016). Convergence to a plausible global maximum was checked by repeatedly refitting the model from randomly chosen starting sets of parameters. All models were fitted in a transformed form to allow only biologically-meaningful values of parameters; that is, the basic reproduction numbers were estimated after transformation with log (*R*_0,*i*_ − 1) to ensure *R*_0,*i*_ > 1.

#### 4.2.3 Model comparison

To compare the best-fitting NiSP or NiDP model and an appropriate model assuming statistical independence (binomial or multinomial), we use the Akaike Information Criterion 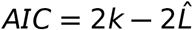, in which 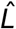 is the log-likelihood of the best-fitting version of each model and *k* is the number of model parameters. This is necessary since these comparisons involve pairs of models that are not nested.

#### 4.2.4 Goodness-of-fit

We use a Monte-Carlo technique to estimate *p*-values for model goodness-of-fit, generating 1, 000, 000 independent sets of samples of total size *N* from the multinomial distribution corresponding to the best-fitting model, calculating the likelihood (Eq. (30)) of each of these synthetic data sets, and recording the proportion with a smaller value of *L* than the value calculated for the data (Sokal and Rohlf, 2012). This was done using the function xmonte() in the R package XNomial (Engels, 2015).

### 4.3 Sources of data and results of model fitting

#### 4.3.1 Numbers of distinct pathogens (NiSP model)

Results of fitting the NiSP model to data from four publications for strains of a single pathogen are presented in Figure 3. Error bars are 95% confidence intervals using exact methods for binomial proportions via binconf() in the R package Hmisc (Harrell Jr et al., 2016). Results for three further data sets concerning different pathogens of a single host (Andersson et al., 2013; Moutailler et al., 2016; Nickbakhsh et al., 2016) are provided as Supplementary Information (Section S2.1).

For convenience the raw data as extracted for use in model fitting are re-tabulated in Table 2. Results of model fitting are summarized in Table 3. We used the value *n* = 102 for the number of distinct strains in López-Villavicencio et al. (2007) following personal communication with the authors; there might be undetected genetic differences due to missing data – which would require a larger value of *n* in our model fitting procedure – but we confirmed that our inferences are unaffected by taking any value of *n* ∈ [100, 200].

**Table 2:**
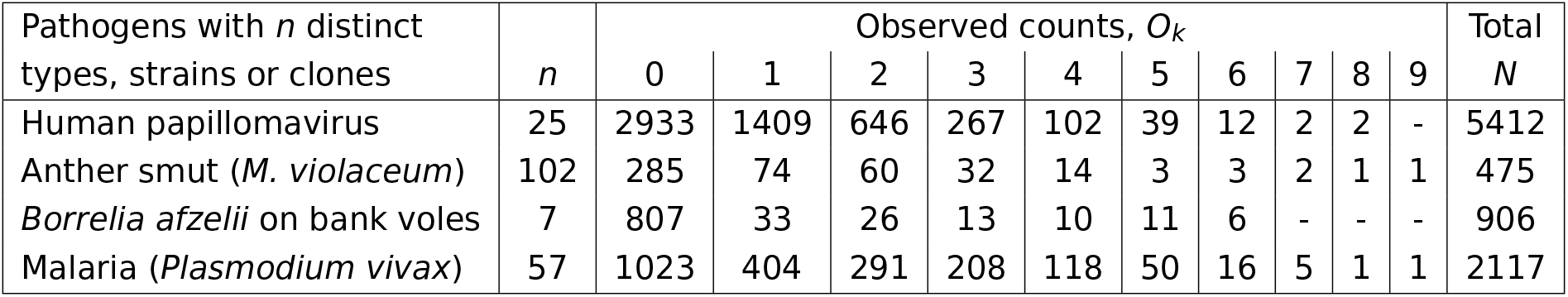
Sources of data for fitting the NiSP model in which pathogen types, clones or strains are assumed to be epidemiologically-interchangeable. The data sets include human papillomavirus (Chaturvedi et al., 2011), anther smut (*M. violaceum*) (López-Villavicencio et al., 2007), *Borrelia afzelii* on bank voles (Andersson et al., 2013), and malaria (*Plasmodium vivax*) (Koepfli et al., 2011).

| Pathogens with $n$ distinct types, strains or clones | $n$ | Observed counts, $O_k$ | | | | | | | | | | Total<br>$N$ |
| --- | --- | --- | --- | --- | --- | --- | --- | --- | --- | --- | --- | --- |
|  |  | 0 | 1 | 2 | 3 | 4 | 5 | 6 | 7 | 8 | 9 |  |
| Human papillomavirus | 25 | 2933 | 1409 | 646 | 267 | 102 | 39 | 12 | 2 | 2 | - | 5412 |
| Anther smut (*M. violaceum*) | 102 | 285 | 74 | 60 | 32 | 14 | 3 | 3 | 2 | 1 | 1 | 475 |
| *Borrelia afzelii* on bank voles | 7 | 807 | 33 | 26 | 13 | 10 | 11 | 6 | - | - | - | 906 |
| Malaria (*Plasmodium vivax*) | 57 | 1023 | 404 | 291 | 208 | 118 | 50 | 16 | 5 | 1 | 1 | 2117 |

**Table 3:**
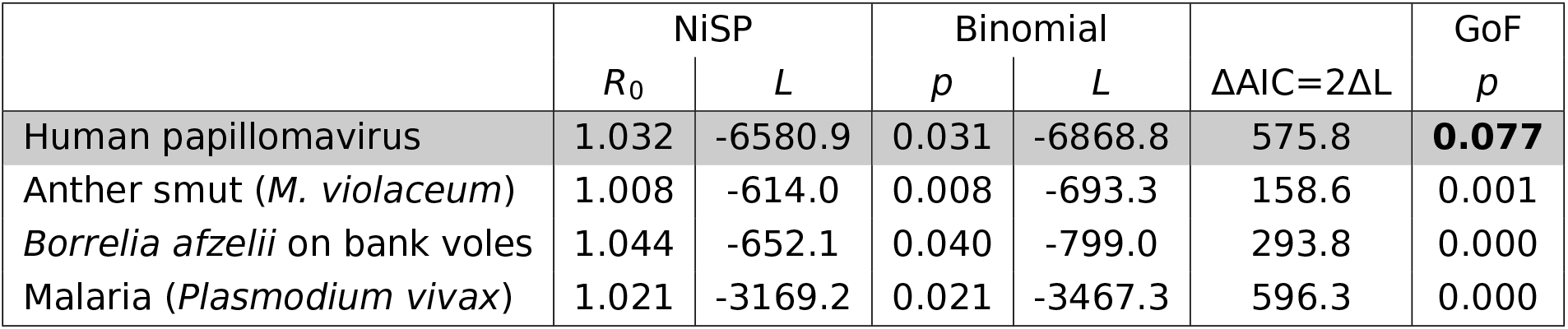
Fitting the NiSP model. The NiSP model was highly supported over the binomial model (ΔAIC ≫ 10) in all cases tested. The final column of the table – GoF – corresponds to the goodness-of-fit test of the NiSP model; values *p* > 0.05 correspond to lack of evidence for failure to fit the data, and so the NiSP model is adequate for the data concerning human papillomavirus (Chaturvedi et al., 2011).

| | NiSP | | Binomial | | $\Delta AIC=2\Delta L$ | GoF<br>$p$ |
| --- | --- | --- | --- | --- | --- | --- |
| | $R_0$ | $L$ | $p$ | $L$ | | |
| Human papillomavirus | 1.032 | -6580.9 | 0.031 | -6868.8 | 575.8 | **0.077** |
| Anther smut (*M. violaceum*) | 1.008 | -614.0 | 0.008 | -693.3 | 158.6 | 0.001 |
| *Borrelia afzelii* on bank voles | 1.044 | -652.1 | 0.040 | -799.0 | 293.8 | 0.000 |
| Malaria (*Plasmodium vivax*) | 1.021 | -3169.2 | 0.021 | -3467.3 | 596.3 | 0.000 |

#### 4.3.2 Combinations of pathogens (NiDP model)

Howard et al. (2001) report results of analyzing 73 data sets concerning multiple *Plasmodium* spp. causing malaria (rows 68–140 of Table 1 in that paper). We re-analyzed the subset of these studies satisfying certain additional constraints as detailed in the main text (see Supplementary Information: Section S2.2 for a full description of how the studies were filtered). This left a final total of 41 data sets taken from 35 distinct papers: 24 data sets considering the three-way interaction between *P. falciparum*, *P. malariae* and *P. vivax* and 17 data sets considering the three-way interaction between *P. falciparum*, *P. malariae* and *P. ovale*.

We used our method based on the NiDP model to test whether any of these data sets were consistent with no interaction between the *Plasmodium* spp. considered (Table 4). We found 15 data sets for which the NiDP model was: i) a better fit than the multinomial model as indicated by ΔAIC ≥ 2; ii) sufficient to explain the data as revealed by our goodness-of-fit test. In these 15 cases our methods therefore support the hypothesis of no interaction. For 11 of these 15 data sets (76, 109, 118, 130, 132, 68, 69, 70, 79, 95, 97, 98, 99, 100, 102) the results as reported in (Howard et al., 2001) instead suggest the strains interact.

**Table 4:** Fitting the NiDP model. Data sets which are consistent with no interaction between the *Plasmodium* spp. considered are highlighted in grey. Such data sets have both *p*-values for the goodness-of-fit test of the NiDP model *p*(GoF) > 0.05, and Δ*AIC* ≥ 2, meaning the NiDP model is adequate. The multinomial model corresponds to the statistical independence hypothesis. Parameters *R*_0,1_ and *R*_0,2_ are associated with *P. falciparum* and *P. malariae*, respectively. Parameter *R*_0,3_ corresponds either to *P. vivax* (upper part of the table, data sets 74–137) or to *P. ovale* (lower part of the table, data sets 68–103). The final column contains a tick whenever at least one association between a pair of pathogens was assessed to be significant in Howard et al. (2001). Red ticks correspond to possible statistical associations that are consistent with our no-interaction model (NiDP), i.e. cases in which our methods lead to results diverging from those reported in Howard et al. (2001).

|  | NiDP |  |  |  |  | Multinomial |  |  |  |  |  | Association(s) in<br>Howard et al. (2001) |
| --- | --- | --- | --- | --- | --- | --- | --- | --- | --- | --- | --- | --- |
| | $R_{0,1}$ | $R_{0,2}$ | $R_{0,3}$ | $L$ | $p(\text{GoF})$ | $p_1$ | $p_2$ | $p_3$ | $L$ | $p(\text{GoF})$ | $\Delta\text{AIC}$ | |
| 74 | 1.764 | 1.256 | 1.004 | -340.7 | 0.000 | 0.468 | 0.220 | 0.004 | -311.0 | 0.000 | -59.3 | ✓ |
| 75 | 1.694 | 1.248 | 1.022 | -194.8 | 0.000 | 0.445 | 0.215 | 0.022 | -177.4 | 0.000 | -34.8 | ✓ |
| **76** | 1.235 | 1.019 | 1.005 | -492.5 | **0.251** | 0.190 | 0.019 | 0.005 | -493.7 | 0.098 | **2.4** | ✓ |
| 82 | 1.776 | 1.165 | 1.108 | -996.0 | 0.000 | 0.463 | 0.147 | 0.101 | -936.3 | 0.000 | -119.2 | ✓ |
| 84 | 1.212 | 1.017 | 1.207 | -684.2 | 0.000 | 0.180 | 0.017 | 0.177 | -660.4 | 0.000 | -47.6 | ✓ |
| 88 | 1.296 | 1.120 | 1.260 | -314.7 | 0.000 | 0.242 | 0.111 | 0.217 | -295.0 | 0.000 | -39.3 | ✓ |
| 106 | 1.818 | 1.146 | 1.055 | -4105.2 | 0.000 | 0.442 | 0.125 | 0.052 | -4296.6 | 0.000 | 382.9 | ✓ |
| 108 | 1.241 | 1.024 | 1.096 | -1147.5 | 0.000 | 0.197 | 0.023 | 0.089 | -1132.1 | 0.721 | -30.9 | ✗ |
| **109** | 1.023 | 1.013 | 1.045 | -359.3 | **0.866** | 0.023 | 0.013 | 0.043 | -361.1 | 0.343 | **3.5** | ✓ |
| 111 | 1.198 | 1.005 | 1.786 | -1929.2 | 0.000 | 0.175 | 0.005 | 0.467 | -1798.8 | 0.000 | -260.7 | ✓ |
| **112** | 1.307 | 1.086 | 1.056 | -119.6 | **0.115** | 0.241 | 0.080 | 0.054 | -116.6 | 0.552 | -6.0 | ✗ |
| 113 | 1.213 | 1.007 | 1.119 | -1324.1 | 0.000 | 0.179 | 0.007 | 0.108 | -1290.8 | 0.000 | -66.6 | ✓ |
| 114 | 1.615 | 1.084 | 1.038 | -1224.4 | 0.000 | 0.392 | 0.080 | 0.037 | -1182.6 | 0.000 | -83.6 | ✓ |
| 116 | 1.780 | 1.124 | 1.100 | -1035.1 | 0.000 | 0.471 | 0.116 | 0.094 | -953.5 | 0.000 | -163.2 | ✓ |
| 117 | 1.072 | 1.000 | 1.268 | -31530.5 | 0.000 | 0.068 | 0.000 | 0.214 | -30958.7 | 0.000 | -1143.5 | ✓ |
| **118** | 1.085 | 1.039 | 1.171 | -225.3 | **0.990** | 0.078 | 0.037 | 0.146 | -227.5 | 0.515 | **4.5** | ✓ |
| 119 | 1.433 | 1.164 | 1.375 | -265.7 | 0.000 | 0.325 | 0.146 | 0.291 | -249.0 | 0.146 | -33.6 | ✓ |
| 123 | 1.016 | 1.055 | 1.098 | -6684.7 | 0.000 | 0.016 | 0.052 | 0.090 | -6623.5 | 0.000 | -122.4 | ✓ |
| 124 | 1.254 | 1.100 | 1.082 | -3600.6 | 0.000 | 0.206 | 0.092 | 0.076 | -3541.3 | 0.017 | -118.7 | ✓ |
| 127 | 1.341 | 1.005 | 1.266 | -1087.4 | 0.000 | 0.265 | 0.005 | 0.219 | -1039.0 | 0.000 | -96.8 | ✓ |
| **130** | 1.013 | 1.002 | 1.350 | -352.7 | **0.978** | 0.013 | 0.002 | 0.259 | -353.7 | 0.636 | **2.0** | ✗ |
| **132** | 1.397 | 1.027 | 1.074 | -591.8 | **0.347** | 0.285 | 0.026 | 0.068 | -594.3 | 0.067 | **4.9** | ✓ |
| 133 | 1.571 | 1.022 | 1.332 | -687.9 | 0.000 | 0.375 | 0.022 | 0.257 | -676.2 | 0.001 | -23.4 | ✓ |
| 137 | 1.196 | 1.005 | 1.130 | -2356.8 | 0.000 | 0.166 | 0.005 | 0.117 | -2309.6 | 0.000 | -94.3 | ✓ |
| **68** | 1.910 | 1.091 | 1.021 | -152.0 | **0.200** | 0.469 | 0.082 | 0.020 | -157.8 | 0.002 | **11.7** | ✓ |
| **69** | 4.827 | 1.443 | 1.036 | -177.2 | **0.822** | 0.796 | 0.310 | 0.035 | -181.4 | 0.121 | **8.5** | ✓ |
| **70** | 4.612 | 1.203 | 1.089 | -239.2 | **0.953** | 0.781 | 0.168 | 0.082 | -247.4 | 0.012 | **16.4** | ✓ |
| 71 | 6.070 | 1.370 | 1.181 | -310.1 | 0.001 | 0.822 | 0.261 | 0.148 | -336.2 | 0.000 | 52.1 | ✗ |
| 77 | 14.275 | 1.383 | 1.142 | -155.3 | 0.032 | 0.944 | 0.286 | 0.127 | -150.4 | 0.931 | -9.9 | ✗ |
| **78** | 4.171 | 1.178 | 1.006 | -166.2 | **0.264** | 0.773 | 0.153 | 0.006 | -163.2 | 0.997 | -5.8 | ✗ |
| **79** | 1.855 | 1.033 | 1.005 | -1260.1 | **0.969** | 0.461 | 0.032 | 0.005 | -1263.4 | 0.224 | **6.6** | ✓ |
| **80** | 1.546 | 1.062 | 1.021 | -715.1 | **0.735** | 0.355 | 0.059 | 0.020 | -715.2 | 0.675 | 0.2 | ✓ |
| **95** | 1.855 | 1.033 | 1.005 | -1260.1 | **0.970** | 0.461 | 0.032 | 0.005 | -1263.4 | 0.224 | **6.6** | ✗ |
| 96 | 1.910 | 1.071 | 1.017 | -240.4 | 0.019 | 0.469 | 0.065 | 0.016 | -248.9 | 0.000 | 17.1 | ✗ |
| **97** | 1.952 | 1.077 | 1.004 | -242.6 | **0.568** | 0.486 | 0.071 | 0.004 | -246.7 | 0.031 | **8.3** | ✓ |
| **98** | 1.662 | 1.014 | 1.018 | -183.7 | **0.373** | 0.396 | 0.013 | 0.018 | -187.0 | 0.030 | **6.6** | ✗ |
| **99** | 1.627 | 1.019 | 1.019 | -133.7 | **0.823** | 0.384 | 0.019 | 0.019 | -135.6 | 0.332 | **3.8** | ✗ |
| **100** | 1.037 | 1.003 | 1.000 | -432.1 | **0.254** | 0.035 | 0.003 | 0.000 | -433.2 | 0.083 | **2.3** | ✓ |
| 101 | 3.590 | 1.269 | 1.063 | -11014.1 | 0.000 | 0.720 | 0.211 | 0.060 | -11392.7 | 0.000 | 757.3 | ✓ |
| **102** | 2.473 | 1.153 | 1.027 | -8188.9 | **0.403** | 0.595 | 0.132 | 0.027 | -8352.0 | 0.000 | **326.2** | ✓ |
| 103 | 1.798 | 1.180 | 1.015 | -7425.7 | 0.000 | 0.437 | 0.150 | 0.015 | -7736.6 | 0.000 | 621.8 | ✓ |

## 5 Data availability

The cross-sectional survey data extracted from previous publications which we have used to test our methodology are tabulated in Table 2 in the main text, and Tables S2 and S4 in the Supplementary Information. These data are available in electronic format as .csv files from the corresponding author upon reasonable request.

## 6 Code availability

Code illustrating all statistical methods is freely available at: https://github.com/nikcunniffe/Coinfection.

## Acknowledgments

This work was initiated during the Multiscale Vectored Plant Viruses Working Group at the National Institute for Mathematical and Biological Synthesis, supported by the National Science Foundation through NSF Award #DBI-1300426, with additional support from The University of Tennessee, Knoxville. This material is based upon research supported by the Thomas Jefferson Fund of the Embassy of France in the United States and the FACE Foundation. We thank S. Alizon, T. Berrett, E. Bussell, V. Calcagno, C. Donnelly, R. Donnelly, T. Giraud, J. Gog, M. López-Villavicencio, T. Obadia, M. Parry, M. Plantegenest, O. Restif, E. Seabloom, J. Shykoff, R. Thompson and C. Trotter for helpful discussions or provision of data.

