## Supporting Information for "Co-infections by non-interacting pathogens are not independent & require new tests of interaction"

### Supporting information for "Co-infections by non-interacting pathogens are not independent and require new tests of interaction"

**Frédéric M. Hamelin, Linda J.S. Allen, Vrushali A. Bokil, Louis J. Gross, Frank M. Hilker, Michael J. Jeger, Carrie A. Manore, Alison G. Power, Megan A. Rúa, Nik J. Cunniffe**

#### Organisation of this document

Supplementary Information S1 ("Mathematical supplements") contains five subsections:

- S1.1. "Formal proof that the proportion of co-infected hosts is larger than the product of the prevalences".
  - A demonstration that $J_{1,2}$ from Eq. (3) in the Main Text is larger than $P = I_1 I_2$ as can be derived from Eq. (1) in the Main Text for all parameters (for sufficiently large $t$).
- S1.2. "Covariance matrix at the endemic equilibrium".
  - A derivation of the covariance matrix for stochastic fluctuations around the endemic equilibrium in the two-pathogen model (leads to Eq. (27) in the main text).
- S1.3. "Comments on the model of May and Nowak (1995)".
  - Details how the often-cited co-infection model of May and Nowak (1995) is not correct.
- S1.4. "Extending the models to accommodate specific clearance".

- Shows how the methods developed in the main text can be extended to accommodate an additional epidemiological parameter: specific clearance (i.e. each pathogen being cleared independently of any other, possibly at a pathogen-specific rate).

- S1.5. "The prevalence of co-infections can be equal to the product of the prevalences of interacting pathogens".
  - Shows how, in the model described in Supplementary Information, Section S1.4, if there is no host natural death but only specific clearance, then the prevalence of co-infections can be equal to the product of the prevalences even when pathogens interact. This complements our main point (non-interaction does not imply independence) by showing that it holds the other way around as well (independence does not imply non-interaction).

Supplementary Information S2 ("Sources of data and side results of model fitting") contains three subsections:

- S2.1. "Additional fitting of the NiSP model".
  - Results of fitting the NiSP model to three additional data sets not considered in the main text. These studies do not concern a single pathogen species, and so the pragmatic assumption of epidemiological interchangeability between pathogens is less justifiable.
- S2.2. "Fitting the NiDP model".
  - Re-tabulates the data sets used by Howard et al. (2001), and describes in full how our criteria to allow a study to be considered led to us fixing on 41 particular studies to analyse.
- S2.3. "Fitting the models with specific clearance".
  - Shows that fitting the NiSP model with specific clearance confirms results found from a model that does not include specific clearance as an additional parameter.

### S1 Mathematical supplements

#### S1.1 Formal proof that the proportion of co-infected hosts is larger than the product of the prevalences

Let $P = I_1 I_2$ and introduce the new variable $Z$, the extent to which the product of prevalences over-estimates the proportion of co-infected hosts,

$$Z = P - J_{1,2} = I_1 I_2 - J_{1,2}. \tag{S1}$$

Differentiating $Z$ and simplifying using Eqs. (2) and (3) in the main text leads to

$$\begin{aligned} \dot{Z} &= I_1 \dot{I}_2 + I_2 \dot{I}_1 - \dot{J}_{1,2}, \\ \dot{Z} &= -(\beta_1 I_1 + \beta_2 I_2 + \mu) Z - \mu P. \end{aligned} \tag{S2}$$

Of interest is the sign of $Z(t)$ as a function of its initial conditions. We assume $R_{0,i} > 1$ and $0 < I_i(0) \le \bar{I}_i$ for $i = 1, 2$. The differential equation for $I_i$ is logistic and therefore, $I_i(t)$ converges monotonically to $\bar{I}_i$, $i = 1, 2$. There exists $\epsilon > 0$ such that $P(t) > \tilde{P} = \bar{I}_1 \bar{I}_2 - \epsilon > 0$ for $t \ge 0$. Suppose $Z(0) > 0$. The term $-\mu P(t) < -\mu \tilde{P} < 0$ ensures that $Z(t)$ decreases to zero in finite time. Let $t_1$ be the first time that $Z(t_1) = 0$. Then $\dot{Z}(t_1) < 0$, so $Z(t)$ eventually becomes negative. To show that $Z(t)$ remains negative, let $Z(t) < 0$ for $t_1 < t < t_2$ and let $t_2$ be the first time that $Z(t_2) = 0$, then $\dot{Z}(t_2) \ge 0$, a contradiction to the fact that $\dot{Z}(t_2) \le -\mu \tilde{P}$. Hence $Z(t)$ remains negative for $t > t_1$. In a similar manner it can be shown that if $Z(0) = 0$ or $Z(0) < 0$, then $Z(t) < 0$ for $t > 0$. In particular, due to the convergence of $I_i(t)$ to $\bar{I}_i$ for $i = 1, 2$, $Z(t)$ converges to the negative limit: $-\mu \bar{I}_1 \bar{I}_2 / (\beta_1 \bar{I}_1 + \beta_2 \bar{I}_2 + \mu)$.

In summary, the fate of $Z$ is to become negative in finite time and to remain negative. This is due to $\mu > 0$. Otherwise for $\mu = 0$, $Z$ would not change sign and would asymptotically converge to zero.

#### S1.2 Covariance matrix at the endemic equilibrium

In the stochastic version of the model, the fluctuations $(\Delta I_1, \Delta I_2) = (I_1 - \bar{I}_1, I_2 - \bar{I}_2)$ about the endemic equilibrium $(\bar{I}_1, \bar{I}_2) = N(1 - 1/R_{0,1}, 1 - 1/R_{0,2})$ can be approximated by the solution of the linear multivariate Fokker-Planck equation,

$$\frac{\partial p(x,t)}{\partial t} = -\sum_{i,j=1}^{2} A_{ij}\frac{\partial (x_j p)}{\partial x_i} + \frac{1}{2}\sum_{i,j=1}^{2} B_{ij}\frac{\partial^2 p}{\partial x_i \partial x_j}\,, \tag{S3}$$

where the vector $x = (x_1, x_2)$ corresponds to $(\Delta I_1, \Delta I_2)$. The steady-state solution of this equation is a Gaussian distribution with mean zero and covariance matrix $\bar{C}$. We will use this multivariate normal distribution to approximate the joint probability density function of the random variables $(I_1, I_2)$ near the endemic equilibrium. Matrix $A = [A_{ij}]$ is the rate of change toward zero and matrix $B = [B_{ij}]$ is the covariance of this process (O'Dea et al., 2018; Van Kampen, 1992). In particular, matrix $A$ is the linearization of the differential equations for $(I_1, I_2)$ (total population size, not proportions) about the endemic equilibrium,

$$A = \begin{bmatrix} \beta_1 - 2\frac{\beta_1}{N}\bar{I}_1 - \mu & 0 \\ 0 & \beta_2 - 2\frac{\beta_2}{N}\bar{I}_1 - \mu \end{bmatrix} = \begin{bmatrix} -\beta_1 + \mu & 0 \\ 0 & -\beta_2 + \mu \end{bmatrix}. \tag{S4}$$

We use the fact that $I_1 = J_1 + J_{1,2}$ and $I_2 = J_2 + J_{1,2}$ and sum the appropriate elements in the covariance matrix $\Sigma$ (Eq. (24) in the main text) to compute the covariance matrix $B$,

$$B = \begin{bmatrix} \bar{F}_1(\bar{J}_\varnothing + \bar{J}_2) + \mu \bar{I}_1 & \mu \bar{J}_{12} \\ \mu \bar{J}_{12} & \bar{F}_2(\bar{J}_\varnothing + \bar{J}_1) + \mu \bar{I}_2 \end{bmatrix} = \begin{bmatrix} 2N\mu\left(1 - \frac{1}{R_{0,1}}\right) & \mu \bar{J}_{12} \\ \mu \bar{J}_{12} & 2N\mu\left(1 - \frac{1}{R_{0,2}}\right) \end{bmatrix} \tag{S5}$$

The expressions in matrices $A$ and $B$ are evaluated at the endemic equilibrium. In particular, $\bar{F}_i = \beta_i \bar{I}_i / N = \beta_i(1 - 1/R_{0,i})$, $i = 1, 2$, and the equilibrium $\bar{J}_{12}$ is found by multiplying Eq. (9) in the main text by $N$:

$$\bar{J}_{12} = \left(\frac{N(\beta_1 + \beta_2)}{\beta_1 + \beta_2 - \mu}\right)\left(1 - \frac{1}{R_{0,1}}\right)\left(1 - \frac{1}{R_{0,2}}\right). \tag{S6}$$

Van Kampen (1992) showed that the covariance matrix $C$ of the Fokker-Planck equation is the solution of the differential equation: $\dot{C} = AC + CA^T + B$. The steady-state covariance matrix is the solution of

$$AC + CA^T = -B\,. \tag{S7}$$

To compute the steady-state covariance matrix for the proportion of the population that is infected, divide the solution of Eq. (S7) by $N^2$. That is, $\bar{C}$ equals

$$\bar{C} = \frac{1}{N^2}\begin{bmatrix} \dfrac{N\mu\left(1-\frac{1}{R_{0,1}}\right)}{\beta_1-\mu} & \dfrac{\mu\bar{J}_{1,2}}{(\beta_1-\mu)+(\beta_2-\mu)} \\ \dfrac{\mu\bar{J}_{1,2}}{(\beta_1-\mu)+(\beta_2-\mu)} & \dfrac{N\mu\left(1-\frac{1}{R_{0,2}}\right)}{\beta_2-\mu} \end{bmatrix} = \begin{bmatrix} \dfrac{1}{NR_{0,1}} & \dfrac{\mu\bar{J}_{1,2}}{N^2[(\beta_1-\mu)+(\beta_2-\mu)]} \\ \dfrac{\mu\bar{J}_{1,2}}{N^2[(\beta_1-\mu)+(\beta_2-\mu)]} & \dfrac{1}{NR_{0,2}} \end{bmatrix}, \tag{S8}$$

where $\bar{J}_{12}$ is defined in Eq. (S6). The steady-state covariance matrix in Eq. (S8) is used to construct confidence ellipses about the endemic equilibrium $(\bar{I}_1/N, \bar{I}_2/N) = (1-1/R_{0,1}, 1-1/R_{0,2})$ (as shown in Fig. 2C in the main text).

Note that the covariance between the prevalences of pathogen 1 and pathogen 2 (the off-diagonal elements in Eq. (S8)) is

$$\bar{C}_{ij} = \text{cov}\left(\frac{I_1}{N}, \frac{I_2}{N}\right) = \frac{\mu\bar{J}_{1,2}}{N^2[(\beta_1-\mu)+(\beta_2-\mu)]} = \frac{(\beta_1+\beta_2)(\beta_1-\mu)(\beta_2-\mu)\mu}{N\beta_1\beta_2(\beta_1+\beta_2-\mu)(\beta_1-\mu+\beta_2-\mu)} \geq 0\,, \tag{S9}$$

(for $i \neq j$) with equality if and only if $\mu = 0$ (assuming $\beta_i > \mu$, $i = 1, 2$).

#### S1.3 Comments on the model of May and Nowak (1995)

May and Nowak (1995) introduced a co-infection model very similar to that presented in the main text, taking

$$\dot{I}_i = I_i[\beta_i(1-I_i) - \mu - \bar{\alpha}_i]\,, \quad \text{with} \quad i = 1, \ldots, n\,, \tag{S10}$$

for $n$ pathogens. The natural mortality rate of the host is $\mu$. The only difference from our model is pathogen-specific mortality. In a single infection, pathogen $i$ induces an additional death rate to the host $\alpha_i$: this is the virulence of pathogen $i$. The induced death rate of co-infected hosts is assumed to be equal to the maximum virulence of the co-infecting pathogens. The pathogens are ranked such that for all $i$, $\alpha_i < \alpha_{i+1}$. Pathogen 1 is the least virulent pathogen and $n$ is the most virulent pathogen. The term $\bar{\alpha}_i$ denotes the average induced death rate of hosts infected by pathogen $i$.

The authors state that the probability that a host is not infected with a pathogen more virulent than $i$ is defined as:

$$p_i = \prod_{j=i+1}^{n} (1 - I_j)\,. \tag{S11}$$

It is important to notice that an underlying assumption of this definition is that the dynamics of the pathogens are independent. But, as we show below, they are not, since the most virulent pathogens influence the dynamics of least virulent pathogens. The coupling term $\bar{\alpha}_i$ is defined as:

$$\bar{\alpha}_i = \alpha_i p_i + \sum_{j=i+1}^{n} \alpha_j I_j p_j\,. \tag{S12}$$

The term $I_j p_j$ represents the probability to be infected by $j$ and uninfected by more virulent pathogens than $j$. Again, this definition implicitly assumes that the dynamics of the pathogens are independent. This seems to contradict the fact that the dynamics are coupled through virulence.

In this section, we check the model given in Eq. (S10) for $n = 2$ pathogens and show that the above definitions do not hold up to mathematical analysis. We consider the same 2-pathogen model as Eq. (3) of the main text, except that we include virulence parameters $\alpha_2 > \alpha_1$. Model (S10) is to be compared with:

$$\begin{aligned}
\dot{J}_1 &= F_1 J_\varnothing - (F_2 + \mu + \alpha_1) J_1\,, \\
\dot{J}_2 &= F_2 J_\varnothing - (F_1 + \mu + \alpha_2) J_2\,, \\
\dot{J}_{1,2} &= F_2 J_1 + F_1 J_2 - (\mu + \max(\alpha_1, \alpha_2)) J_{1,2}\,, \\
&= F_2 J_1 + F_1 J_2 - (\mu + \alpha_2) J_{1,2}\,,
\end{aligned} \tag{S13}$$

where $J_\varnothing = 1 - J_1 - J_2 - J_{1,2}$.

Since model (S10) and model (S13) share the same biological assumptions and the same mathematical formalism, they should be equivalent (for $n = 2$ pathogens). Let $I_1 = J_1 + J_{1,2}$ and $I_2 = J_2 + J_{1,2}$. Model (S13) is equivalent to

$$\begin{aligned}
\dot{I}_1 &= \beta_1 I_1 (1 - I_1) - (\mu + \alpha_1^\star) I_1\,, \\
\dot{I}_2 &= \beta_2 I_2 (1 - I_2) - (\mu + \alpha_2) I_2\,, \\
\dot{J}_{1,2} &= \beta_1 I_1 (I_2 - J_{1,2}) + \beta_2 I_2 (I_1 - J_{1,2}) - (\mu + \alpha_2) J_{1,2}\,,
\end{aligned} \tag{S14}$$

where

$$\alpha_1^\star = \alpha_1 \left(1 - \frac{J_{1,2}}{I_1}\right) + \alpha_2 \frac{J_{1,2}}{I_1} .$$

Eq. (S11) yields $p_1 = 1 - I_2$ and $p_2 = 1$. Eq. (S12) yields

$$\bar{\alpha}_1 = \alpha_1(1 - I_2) + \alpha_2 I_2 .$$

For model (S10) and model (S13) to coincide, one must have $\alpha_1^\star = \bar{\alpha}_1$, i.e. $J_{1,2} = I_1 I_2$. Proceeding as in Supplementary Information, Section S1.1, let $P = I_1 I_2$ and $Z = P - J_{1,2}$. We have

$$\dot{Z} = -(\beta_1 I_1 + \beta_2 I_2 + \mu + \alpha_2)Z - (\mu + \alpha_1^\star)P .$$

Assuming $P(t) > \tilde{P} > 0$ for $t > 0$, it can be shown that $Z(t)$ becomes negative and stays negative, implying for some time $t_0$ and $t > t_0$, $J_{1,2}(t) > P(t) = I_1(t)I_2(t)$. Therefore, $\alpha_1^\star \neq \bar{\alpha}_1$. Hence, model (S10) and model (S13) are not equivalent, as they should be, if model (S10) is correct.

#### S1.4 Extending the models to accommodate specific clearance

##### S1.4.1 Two-pathogen model

Introducing a pathogen-specific clearance rate $\gamma_i$, Eq. (1) of the main text is replaced by

$$\dot{I}_i = \beta_i I_i (1 - I_i) - (\gamma_i + \mu) I_i \tag{S15}$$

and Eq. (3) by

$$\begin{aligned}
\dot{J}_1 &= F_1 J_\varnothing - (F_2 + \gamma_1 + \mu) J_1 + \gamma_2 J_{1,2} , \\
\dot{J}_2 &= F_2 J_\varnothing - (F_1 + \gamma_2 + \mu) J_2 + \gamma_1 J_{1,2} , \\
\dot{J}_{1,2} &= F_2 J_1 + F_1 J_2 - (\gamma_1 + \gamma_2 + \mu) J_{1,2} ,
\end{aligned} \tag{S16}$$

where the definition of $F_i$ is the same as in Eq. (2). The parameter $\mu$ is unchanged: this is the natural death rate of the host.

After inclusion of pathogen-specific clearance rates, Eq. (7) is replaced by

$$\dot{J}_{\varnothing} = \mu(J_1+J_2+J_{1,2}) - (F_1+F_2)J_{\varnothing} + \gamma_1 J_1 + \gamma_2 J_2 = \mu(1-J_{\varnothing}) - (F_1+F_2)J_{\varnothing} + \gamma_1 J_1 + \gamma_2 J_2 \,. \quad \text{(S17)}$$

and the basic reproduction number is

$$R_{0,i} = \frac{\beta_i}{\gamma_i + \mu} \,. \quad \text{(S18)}$$

Also, Eq. (8) is replaced by

$$\dot{J}_{1,2} = \beta_2 I_2 (I_1 - J_{1,2}) + \beta_1 I_1 (I_2 - J_{1,2}) - (\gamma_1 + \gamma_2 + \mu) J_{1,2} \,. \quad \text{(S19)}$$

Eq. (9) is unchanged as the relative deviation from statistical independence is unaffected by the specific clearance rates $\gamma_i$.

Finally, Eq. (S2) is replaced by

$$\dot{Z} = -(\beta_1 I_1 + \beta_2 I_2 + \gamma_1 + \gamma_2 + \mu) Z - \mu P. \quad \text{(S20)}$$

where $Z(t)$ converges to the negative limit: $-\mu \bar{I}_1 \bar{I}_2 / (\beta_1 \bar{I}_1 + \beta_2 \bar{I}_2 + \gamma_1 + \gamma_2 + \mu)$. Again, the fate of $Z$ is to become negative and to remain negative provided $\mu > 0$.

##### S1.4.2 Analysis of the $n$-pathogen model

Introducing the notation for the set of hosts infected by one additional pathogen $\Lambda_i = \Gamma \cup \{i\}$ (for $i \notin \Gamma$), Eq. (5) in the main text becomes

$$\dot{J}_{\Gamma} = \sum_{i \in \Gamma} F_i J_{\Omega_i} - \left( \sum_{i \notin \Gamma} F_i + \sum_{i \in \Gamma} \gamma_i + \mu \right) J_{\Gamma} + \sum_{i \notin \Gamma} \gamma_i J_{\Lambda_i} \,. \quad \text{(S21)}$$

with $F_i$ the same as in Eq. (6). The final term in Eq. (S21) tracks the inflow due to hosts with one additional infection that clear a single infection. This final term is omitted in the single case in which $\Gamma$ corresponds to infection by all pathogens. Also, the updated version of Eq. (10) for $\dot{J}_{\varnothing}$ with pathogen-specific clearance rates is

$$\dot{J}_{\varnothing} = \mu(1 - J_{\varnothing}) - \left( \sum_{i=1}^{n} F_i \right) J_{\varnothing} + \sum_{i=1}^{n} \gamma_i J_i \,. \quad \text{(S22)}$$

**Equilibrium analysis.** The equilibrium equations with pathogen-specific clearance rates are

$$0 = \sum_{i\in\Gamma} \bar{F}_i \bar{J}_{\Omega_i} - \left( \sum_{i\notin\Gamma} \bar{F}_i + \sum_{i\in\Gamma} \gamma_i + \mu \right) \bar{J}_\Gamma + \sum_{i\notin\Gamma} \gamma_i \bar{J}_{\Lambda_i}\,, \tag{S23}$$

and

$$0 = \sum_{i\in\Gamma} (\beta_i - (\gamma_i + \mu)) \bar{J}_{\Omega_i} - \left( \sum_{i\notin\Gamma} (\beta_i - (\gamma_i + \mu)) + \sum_{i\in\Gamma} \gamma_i + \mu \right) \bar{J}_\Gamma + \sum_{i\notin\Gamma} \gamma_i \bar{J}_{\Lambda_i}\,. \tag{S24}$$

with

$$\bar{F}_i = \beta_i \bar{I}_i = \beta_i \left( 1 - \frac{\gamma_i + \mu}{\beta_i} \right) = \beta_i - (\gamma_i + \mu). \tag{S25}$$

(replacing Eqs. (11-12-13)).

To fit the models to data, it would be necessary to scale by the rate of host natural death $\mu$ in Eq. (S24), leading to

$$0 = \sum_{i\in\Gamma} \big(\hat{\beta}_i - (\hat{\gamma}_i + 1)\big) \bar{J}_{\Omega_i} - \left( \sum_{i\notin\Gamma} \big(\hat{\beta}_i - (\hat{\gamma}_i + 1)\big) + \sum_{i\in\Gamma} \hat{\gamma}_i + 1 \right) \bar{J}_\Gamma + \sum_{i\notin\Gamma} \hat{\gamma}_i \bar{J}_{\Lambda_i}, \tag{S26}$$

and so consider infection ($\hat{\beta}_i = \beta_i/\mu$) and specific clearance ($\hat{\gamma}_i = \gamma_i/\mu$) rates measured relative to the rate of host natural death. With the scaled force of infection at equilibrium

$$\hat{F}_i = \hat{\beta}_i - (\hat{\gamma}_i + 1), \tag{S27}$$

then Eq. (S26) can be written as

$$0 = \sum_{i\in\Gamma} \hat{F}_i \bar{J}_{\Omega_i} - \left( \sum_{i\notin\Gamma} \hat{F}_i + \sum_{i\in\Gamma} \hat{\gamma}_i + 1 \right) \bar{J}_\Gamma + \sum_{i\notin\Gamma} \hat{\gamma}_i \bar{J}_{\Lambda_i}\,. \tag{S28}$$

Given the values of $\hat{\beta}_i$ and $\hat{\gamma}_i$, the $2^n-1$ linear equations corresponding to Eq. (S28) can be solved simultaneously with the corresponding equation for the equilibrium density of uninfected hosts (i.e. the scaled version of Eq. (S22)):

$$-1 = -\left( \sum_{i=1}^{n} \hat{F}_i + 1 \right) \bar{J}_\varnothing + \sum_{i=1}^{n} \hat{\gamma}_i \bar{J}_i\,. \tag{S29}$$

to find all $2^n$ equilibrium prevalences predicted by the $n$-pathogen model. However, since the recursive solution presented in the main text (Eq. (16)) is no longer available, the system must be solved using (standard) numerical methods for linear systems of equations.

**Worked example.** When $n = 3$ there is a total of $2^3 = 8$ classes of hosts, uninfected ($J_\varnothing$), singly-infected ($J_1$, $J_2$ and $J_3$), doubly-infected ($J_{1,2}$, $J_{1,3}$ and $J_{2,3}$) and triply-infected ($J_{1,2,3}$). The equilibrium prevalences can be concatenated into a single vector, given here in lexicographical order

$$\mathbf{v} = \left[\bar{J}_\varnothing, \bar{J}_1, \bar{J}_2, \bar{J}_3, \bar{J}_{1,2}, \bar{J}_{1,3}, \bar{J}_{2,3}, \bar{J}_{1,2,3}\right]^T . \tag{S30}$$

If we define $\mathbf{b}$ as

$$\mathbf{b} = [-1, 0, 0, 0, 0, 0, 0, 0]^T , \tag{S31}$$

then Eq. (S28) and (S29) are equivalent to the system of 8 linear equations

$$H\mathbf{v} = \mathbf{b}, \tag{S32}$$

in which matrix $H$ equals

$$\begin{bmatrix}
-(\hat{F}_1+\hat{F}_2+\hat{F}_3+1) & \hat{\gamma}_1 & \hat{\gamma}_2 & \hat{\gamma}_3 & 0 & 0 & 0 & 0 \\
\hat{F}_1 & -(\hat{F}_2+\hat{F}_3+\hat{\gamma}_1+1) & 0 & 0 & \hat{\gamma}_2 & \hat{\gamma}_3 & 0 & 0 \\
\hat{F}_2 & 0 & -(\hat{F}_1+\hat{F}_3+\hat{\gamma}_2+1) & 0 & \hat{\gamma}_1 & 0 & \hat{\gamma}_3 & 0 \\
\hat{F}_3 & 0 & 0 & -(\hat{F}_1+\hat{F}_2+\hat{\gamma}_3+1) & 0 & \hat{\gamma}_1 & \hat{\gamma}_2 & 0 \\
0 & \hat{F}_2 & \hat{F}_1 & 0 & -(\hat{F}_3+\hat{\gamma}_1+\hat{\gamma}_2+1) & 0 & 0 & \hat{\gamma}_3 \\
0 & \hat{F}_3 & 0 & \hat{F}_1 & 0 & -(\hat{F}_2+\hat{\gamma}_1+\hat{\gamma}_3+1) & 0 & \hat{\gamma}_2 \\
0 & 0 & \hat{F}_3 & \hat{F}_2 & 0 & 0 & -(\hat{F}_1+\hat{\gamma}_2+\hat{\gamma}_3+1) & \hat{\gamma}_1 \\
0 & 0 & 0 & 0 & \hat{F}_3 & \hat{F}_2 & \hat{F}_1 & -(\hat{\gamma}_1+\hat{\gamma}_2+\hat{\gamma}_3+1)
\end{bmatrix}.$$

The equilibrium prevalence of hosts infected by any combination of pathogens can then be obtained by solving Eq. (S32) for $\mathbf{v}$.

**Proof that there is always a unique equilibrium.** For the case $n = 3$ pathogens, the matrix $-H$ has off-diagonal entries that are non-positive and diagonal entries that are strictly positive. In addition, the absolute value of each diagonal entry is strictly greater than the absolute value of the sum of all of the other entries in that column. These properties of $-H$ make it a non-singular M-matrix. (Properties of an M-matrix are given in (Plemmons, 1977).) As a consequence of these properties, $-H^{-1}$ exists and is a non-negative matrix from which it follows that the solution $\mathbf{v}$ in Eq. (S32) is non-negative and can be expressed as

$$\mathbf{v} = H^{-1}\mathbf{b}. \tag{S33}$$

Generalizing to the case of $n$ pathogens, it can be verified that matrix $-H$ in Eq. (S32) will still have the same properties, making it a non-singular M-matrix and therefore, the equilibrium $\mathbf{v}$ is the unique non-negative solution given by Eq. (S33).

##### S1.4.3 Relationship between the NiDP and multinomial models

In this subsection, we show that the equilibrium prevalences in the NiDP model with $\mu = 0$ are equal to the expectations under statistical independence, i.e.,

$$\bar{J}_\Gamma = \prod_{i\in\Gamma} \bar{I}_i \prod_{j\notin\Gamma} (1-\bar{I}_j)\,, \tag{S34}$$

where $\bar{I}_i = 1-\gamma_i/\beta_i$ for all $i \in \{1,2,...,n\}$. In other words, when there is no host natural death (at the timescale of an infection), the probability to be infected by a set of pathogens $\Gamma$ follows a multinomial distribution with parameters $n$ (the number of distinct pathogens) and $p_i = \bar{I}_i$ for all $i \in \{1,2,...,n\}$.

In the specific case $\mu = 0$, Eq. (S23) becomes

$$0 = \sum_{i\in\Gamma} \bar{F}_i \bar{J}_{\Omega_i} - \left( \sum_{i\notin\Gamma} \bar{F}_i + \sum_{i\in\Gamma} \gamma_i \right) \bar{J}_\Gamma + \sum_{i\notin\Gamma} \gamma_i \bar{J}_{\Lambda_i}\,, \tag{S35}$$

with $\bar{F}_i = \beta_i - \gamma_i$. Eq. (S34) implies

$$\bar{J}_{\Omega_i} = \bar{J}_\Gamma \frac{1-\bar{I}_i}{\bar{I}_i}\,, \quad \text{and} \quad \bar{J}_{\Lambda_i} = \bar{J}_\Gamma \frac{\bar{I}_i}{1-\bar{I}_i}\,. \tag{S36}$$

Substituting the values in Eq. (S36) into the right side of Eq. (S35),

$$\sum_{i\in\Gamma} \left( \bar{F}_i \frac{1-\bar{I}_i}{\bar{I}_i} \right) - \left( \sum_{i\notin\Gamma} \bar{F}_i + \sum_{i\in\Gamma} \gamma_i \right) + \sum_{i\notin\Gamma} \gamma_i \frac{\bar{I}_i}{1-\bar{I}_i}$$

and simplifying leads to

$$\sum_{i\in\Gamma} \left( (\beta_i - \gamma_i) \frac{\gamma_i}{\beta_i - \gamma_i} \right) - \sum_{i\notin\Gamma} (\beta_i - \gamma_i) + \sum_{i\in\Gamma} \gamma_i + \sum_{i\notin\Gamma} \gamma_i \frac{\beta_i - \gamma_i}{\gamma_i} = 0.$$

Therefore, the values in Eq. (S34) are equilibrium values.

Similarly, in the specific case $\mu = 0$, the equilibrium value for $J_\varnothing > 0$ in Eq. (S22) satisfies

$$0 = \left(\sum_{i=1}^{n} \bar{F}_i\right)\bar{J}_\varnothing + \sum_{i=1}^{n} \gamma_i \bar{J}_i\,. \tag{S37}$$

Applying Eq. (S36) and dividing by $\bar{J}_\varnothing$ in the right side of the preceding equation yields

$$\begin{aligned}\left(\sum_{i=1}^{n} \bar{F}_i\right) + \sum_{i=1}^{n} \gamma_i \frac{\bar{I}_i}{1-\bar{I}_i} &= \left(\sum_{i=1}^{n} (\beta_i - \gamma_i)\right) + \sum_{i=1}^{n} \gamma_i \frac{\beta_i - \gamma_i}{\gamma_i}\,, \\ &= 0\,.\end{aligned}$$

Hence, Eq. (S34) is the equilibrium solution of the NiDP model in the specific case $\mu = 0$.

##### S1.4.4 Relationship between the NiDP and NiSP models

Assuming all pathogens are interchangeable, Eq. (17) of the main text can be replaced by

$$0 = |\Gamma|\hat{F}\bar{J}_{\Omega_i} - \big((n-|\Gamma|)\hat{F} + |\Gamma|\hat{\gamma} + 1\big)\bar{J}_\Gamma + (n-|\Gamma|)\hat{\gamma}\bar{J}_{\Lambda_i}, \tag{S38}$$

in which

$$\hat{F} = \hat{\beta} - (\hat{\gamma} + 1)\,. \tag{S39}$$

For $1 \le k < n$, substituting Eq. (19) into Eq. (S38) leads to

$$0 = k\hat{F}\frac{\bar{M}_{k-1}}{C^n_{k-1}} - \big((n-k)\hat{F} + k\hat{\gamma} + 1\big)\frac{\bar{M}_k}{C^n_k} + (n-k)\hat{\gamma}\frac{\bar{M}_{k+1}}{C^n_{k+1}}. \tag{S40}$$

Noting that

$$\frac{C^n_{k+1}}{C^n_{k-1}} = \frac{(n-k+1)(n-k)}{(k+1)k} \quad \text{and} \quad \frac{C^n_{k+1}}{C^n_k} = \frac{n-k}{k+1},$$

it follows that

$$0 = (n-k+1)\hat{F}\bar{M}_{k-1} - \big((n-k)\hat{F} + k\hat{\gamma} + 1\big)\bar{M}_k + (k+1)\hat{\gamma}\bar{M}_{k+1}, \tag{S41}$$

which holds for $1 \le k < n$ (i.e. there is a total of $n-1$ such equations).

When $k = n$ the analogue of Eq. (S40) is

$$0 = n\hat{F}\frac{\bar{M}_{n-1}}{C^n_{n-1}} - (n\hat{\gamma} + 1)\frac{\bar{M}_n}{C^n_n},$$

and so, since $C^n_{n-1} = n$ and $C^n_n = 1$, it follows that

$$0 = \hat{F}\bar{M}_{n-1} - (n\hat{\gamma} + 1)\bar{M}_n. \tag{S42}$$

When $k = 0$ the analogue of Eqn. (S40) obtained by substituting Eq. (20) into Eq. (S29), is

$$-\left(n\hat{F} + 1\right)\frac{\bar{M}_0}{C^n_0} + n\hat{\gamma}\frac{\bar{M}_1}{C^n_1} = -1, \tag{S43}$$

and so, since $C^n_1 = n$ and $C^n_0 = 1$, it follows that

$$-\left(n\hat{F} + 1\right)\bar{M}_0 + \hat{\gamma}\bar{M}_1 = -1\,. \tag{S44}$$

Taken together, Eqs. (S41-S42-S44) constitute a system of $n + 1$ linear equations that fix the equilibrium prevalences of hosts infected by any number of distinct pathogens in the NiSP model.

**Worked example.** When $n = 3$ there is a total of $n + 1 = 4$ classes of host: uninfected ($M_0$), singly-infected ($M_1$), doubly-infected ($M_2$) and triply-infected ($M_3$). The equilibrium prevalences can be concatenated into a single vector

$$\mathbf{v} = \left[\bar{M}_0, \bar{M}_1, \bar{M}_2, \bar{M}_3\right]^T\,. \tag{S45}$$

If we define $\mathbf{b}$ as

$$\mathbf{b} = [-1, 0, 0, 0]^T\,, \tag{S46}$$

then Eq. (S41-S42-S44) are equivalent to the system of 4 linear equations

$$H\mathbf{v} = \mathbf{b}, \tag{S47}$$

in which matrix $H$ equals

$$\begin{bmatrix} -(3F+1) & \hat{\gamma} & 0 & 0 \\ 3\hat{F} & -(2\hat{F}+\hat{\gamma}+1) & 2\hat{\gamma} & 0 \\ 0 & 2\hat{F} & -(\hat{F}+2\hat{\gamma}+1) & 3\hat{\gamma} \\ 0 & 0 & \hat{F} & -(3\hat{\gamma}+1) \end{bmatrix}. \tag{S48}$$

The equilibrium prevalences of hosts infected by any number of distinct pathogens can then be obtained by solving Eq. (S47).

##### S1.4.5 Relationship between the NiSP and binomial models

In this subsection, we show that the equilibrium prevalences in the NiSP model with $\mu = 0$ are equal to the expectations under statistical independence, i.e.,

$$\bar{M}_k = C_k^n \bar{I}^k (1-\bar{I})^{n-k}\,, \tag{S49}$$

in which $\bar{I} = 1-\gamma/\beta$. In other words, the probability to be infected by $k$ epidemiologically-interchangeable pathogens follows a binomial distribution with parameters $n$ (the number of pathogens considered) and $p = \bar{I}$.

In the specific case $\mu = 0$, Eq. (S38) becomes

$$0 = |\Gamma|\bar{F}\bar{J}_{\Omega_i} - \big((n-|\Gamma|)\bar{F} + |\Gamma|\gamma\big)\bar{J}_{\Gamma} + (n-|\Gamma|)\gamma\bar{J}_{\Lambda_i}\,, \tag{S50}$$

in which $\bar{F} = \beta - \gamma$. Eq. (S41) becomes

$$0 = (n-k+1)\bar{F}\bar{M}_{k-1} - \big((n-k)\bar{F} + k\gamma\big)\bar{M}_k + (k+1)\gamma\bar{M}_{k+1}. \tag{S51}$$

Eq. (S49) implies

$$\bar{M}_{k-1} = \frac{C_{k-1}^n}{C_k^n}\frac{1-\bar{I}}{\bar{I}}\bar{M}_k = \frac{k}{n-k+1}\frac{1-\bar{I}}{\bar{I}}\,, \quad \text{and} \quad \bar{M}_{k+1} = \frac{C_{k+1}^n}{C_k^n}\frac{\bar{I}}{1-\bar{I}}\bar{M}_k = \frac{n-k}{k+1}\frac{\bar{I}}{1-\bar{I}}\bar{M}_k\,. \tag{S52}$$

Substituting the values in Eq. (S52) into the right side of Eq. (S51)

$$(n-k+1)\bar{F}\frac{k}{n-k+1}\frac{1-\bar{I}}{\bar{I}} - \big((n-k)\bar{F} + k\gamma\big) + (k+1)\gamma\frac{n-k}{k+1}\frac{\bar{I}}{1-\bar{I}}$$

and simplifying leads to:

$$\bar{F}k\frac{1-\bar{I}}{\bar{I}}-(n-k)\bar{F}-k\gamma+\gamma(n-k)\frac{\bar{I}}{1-\bar{I}}=k\gamma-(n-k)\bar{F}-k\gamma+(n-k)\bar{F}=0\,.$$

Therefore, the values in Eq. (S49) are equilibrium values.

Similarly, in the specific case $\mu=0$, Eq. (S42) becomes

$$0=\bar{F}\bar{M}_{n-1}-(n\gamma)\,\bar{M}_n\,,$$

and substituting the values in Eq. (S52) leads to

$$\bar{F}n\frac{1-\bar{I}}{\bar{I}}-n\gamma=n\gamma-n\gamma=0\,.$$

Lastly, in the specific case $\mu=0$, Eq. (S44) becomes

$$0=-\left(n\bar{F}\right)\bar{M}_0+\gamma\bar{M}_1\,,$$

and substituting the values in Eq. (S52) leads to

$$-n\bar{F}+\gamma n\frac{\bar{I}}{1-\bar{I}}=-n(\beta-\gamma)+n(\beta-\gamma)=0\,.$$

Hence, Eq. (S49) is the equilibrium solution of the NiSP model in the specific case $\mu=0$.

##### S1.4.6 Stochastic models

**Continuous-time Markov chain.** The continuous-time Markov chain model with pathogen-specific clearance rates has four additional events defined in Table S1.

| Event number | Event | Rate | Change(s) to state variable(s) ($\Delta X$) |
|---|---|---|---|
| 8 | Specific clearance of pathogen 1 from host singly-infected by pathogen 2 | $\gamma_1 J_1 \Delta t + o(\Delta t)$ | $J_1 \to J_1 - 1$<br>$J_\varnothing \to J_\varnothing + 1$ |
| 9 | Specific clearance of pathogen 2 from host singly-infected by pathogen 1 | $\gamma_2 J_2 \Delta t + o(\Delta t)$ | $J_2 \to J_2 - 1$<br>$J_\varnothing \to J_\varnothing + 1$ |
| 10 | Specific clearance of pathogen 1 from co-infected host | $\gamma_1 J_{1,2} \Delta t + o(\Delta t)$ | $J_{1,2} \to J_{1,2} - 1$<br>$J_2 \to J_2 + 1$ |
| 11 | Specific clearance of pathogen 2 from co-infected host | $\gamma_2 J_{1,2} \Delta t + o(\Delta t)$ | $J_{1,2} \to J_{1,2} - 1$<br>$J_1 \to J_1 + 1$ |

Table S1: Additional transitions in the two-pathogen stochastic models.

**Stochastic differential equations.** Let $\mathrm{d}J = \tilde{f}\mathrm{d}t$ be the unscaled version of the deterministic model as specified in Eq. (S16-S17). The extension of matrix $\Sigma$ in Eq. (24) of the main text, to include pathogen specific clearance is

$$\begin{bmatrix} \mu(N-J_\varnothing)+(F_1+F_2)J_\varnothing+\gamma_1 J_1+\gamma_2 J_2 & -F_1 J_\varnothing-(\mu+\gamma_1)J_1 & -F_2 J_\varnothing-(\mu+\gamma_2)J_2 & -\mu J_{1,2} \\ -F_1 J_\varnothing-(\mu+\gamma_1)J_1 & F_1 J_\varnothing+(F_2+\gamma_1+\mu)J_1+\gamma_2 J_{1,2} & 0 & -F_2 J_1-\gamma_2 J_{1,2} \\ -F_2 J_\varnothing-(\mu+\gamma_2)J_2 & 0 & F_2 J_\varnothing+(F_1+\gamma_2+\mu)J_2+\gamma_1 J_{1,2} & -F_1 J_2-\gamma_1 J_{1,2} \\ -\mu J_{1,2} & -F_2 J_1-\gamma_2 J_{1,2} & -F_1 J_2-\gamma_1 J_{1,2} & F_2 J_1+F_1 J_2+(\mu+\gamma_1+\gamma_2)J_{1,2} \end{bmatrix}, \quad \text{(S53)}$$

where $N-J_\varnothing = J_1+J_2+J_{1,2}$ and $N$ is constant.

The new matrix $G$ has dimension $4 \times 11$ due to the four additional events in Table S1, (see Eq. (26)),

$$\begin{aligned}
\mathrm{d}J_\varnothing &= \tilde{f}_0\mathrm{d}t - \sqrt{F_1 J_\varnothing}\,\mathrm{d}W_1 - \sqrt{F_2 J_\varnothing}\,\mathrm{d}W_2 + \sqrt{\mu J_1}\,\mathrm{d}W_5 + \sqrt{\mu J_2}\,\mathrm{d}W_6 + \sqrt{\mu J_{1,2}}\,\mathrm{d}W_7 \\
&\quad + \sqrt{\gamma_1 J_1}\,\mathrm{d}W_8 + \sqrt{\gamma_2 J_2}\,\mathrm{d}W_9\,, \\
\mathrm{d}J_1 &= \tilde{f}_1\mathrm{d}t + \sqrt{F_1 J_\varnothing}\,\mathrm{d}W_1 - \sqrt{F_2 J_1}\,\mathrm{d}W_4 - \sqrt{\mu J_1}\,\mathrm{d}W_5 - \sqrt{\gamma_1 J_1}\,\mathrm{d}W_8 + \sqrt{\gamma_2 J_{1,2}}\,\mathrm{d}W_{11}\,, \quad \text{(S54)} \\
\mathrm{d}J_2 &= \tilde{f}_2\mathrm{d}t + \sqrt{F_2 J_\varnothing}\,\mathrm{d}W_2 - \sqrt{F_1 J_2}\,\mathrm{d}W_3 - \sqrt{\mu J_2}\,\mathrm{d}W_6 - \sqrt{\gamma_2 J_2}\,\mathrm{d}W_9 + \sqrt{\gamma_1 J_{1,2}}\,\mathrm{d}W_{10}\,, \\
\mathrm{d}J_{1,2} &= \tilde{f}_{1,2}\mathrm{d}t + \sqrt{F_1 J_2}\,\mathrm{d}W_3 + \sqrt{F_2 J_1}\,\mathrm{d}W_4 - \sqrt{\mu J_{1,2}}\,\mathrm{d}W_7 - \sqrt{\gamma_1 J_{1,2}}\,\mathrm{d}W_{10} - \sqrt{\gamma_2 J_{1,2}}\,\mathrm{d}W_{11}\,.
\end{aligned}$$

**Covariance matrix at the endemic equilibrium.** The new matrices $A$ and $B$ (Eq. (S4)) are

$$A = \begin{bmatrix} -\beta_1+\gamma_1+\mu & 0 \\ 0 & -\beta_2+\gamma_2+\mu \end{bmatrix}$$

and

$$B = \begin{bmatrix} 2N(\gamma_1+\mu)\left(1-\dfrac{1}{R_{0,1}}\right) & \mu\bar{J}_{12} \\ \mu\bar{J}_{12} & 2N(\gamma_2+\mu)\left(1-\dfrac{1}{R_{0,2}}\right) \end{bmatrix}. \quad \text{(S55)}$$

The new steady-state covariance matrix $\bar{C}$ (Eq. (S8)) is

$$\bar{C} = \begin{bmatrix} \dfrac{1}{NR_{0,1}} & \dfrac{\mu\bar{J}_{1,2}}{N^2[\beta_1-(\gamma_1+\mu)+\beta_2-(\gamma_2+\mu)]} \\ \dfrac{\mu\bar{J}_{1,2}}{N^2[\beta_1-(\gamma_1+\mu)+\beta_2-(\gamma_2+\mu)]} & \dfrac{1}{NR_{0,2}} \end{bmatrix} \quad \text{(S56)}$$

where $\bar{J}_{12}$ is defined in Eq. (9) of the main text.

The covariance between pathogen 1 and pathogen 2 prevalences (the off-diagonal elements in Eq. (S56)) is

$$\begin{aligned}\bar{C}_{ij} &= \mathrm{cov}\left(\frac{I_1}{N},\frac{I_2}{N}\right)=\frac{\mu\bar{J}_{1,2}}{N^2[\beta_1-(\gamma_1+\mu)+\beta_2-(\gamma_2+\mu)]}\\ &= \frac{(\beta_1+\beta_2)(\beta_1-\gamma_1-\mu)(\beta_2-\gamma_2-\mu)\mu}{N\beta_1\beta_2(\beta_1+\beta_2-\mu)(\beta_1-\gamma_1-\mu+\beta_2-\gamma_2-\mu)}\geq 0\,,\end{aligned}$$

(for $i \neq j$) with equality if and only if $\mu = 0$ again (assuming $\beta_i > \gamma_i + \mu$, $i = 1,2$); in the latter case, the deviation from statistical independence is zero (Eq. (9)).

In the special case that $\beta_1 = \beta_2 = \beta$ and $\gamma_1 = \gamma_2 = \gamma$,

$$\frac{\partial \bar{C}_{ii}}{\partial \gamma} = -\frac{\mu}{\beta N(2\beta-\mu)} \leq 0\,,$$

meaning that the positive covariance decreases as $\gamma$ increases (unless $\mu = 0$), as expected.

#### S1.5 The prevalence of co-infections can be equal to the product of the prevalences of interacting pathogens

We consider the same two-pathogen model as Eq. (S16), except we let $\mu = 0$. However, we include two interaction parameters $\sigma_1, \sigma_2 > -1$, such that the forces of infection of both pathogens are

$$F_1 = \beta_1(J_1 + (1+\sigma_1)J_{1,2})\,,\quad F_2 = \beta_2(J_2 + (1+\sigma_2)J_{1,2})\,. \tag{S57}$$

If $\sigma_i < 0$ (resp. $> 0$), then transmission of pathogen $i$ from a co-infected host is lower (resp. greater) than from singly infected hosts ($i = 1,2$). With these assumptions, the model is

$$\begin{aligned}\dot{J}_1 &= F_1 J_\varnothing - (F_2+\gamma_1)J_1 + \gamma_2 J_{1,2}\,,\\ \dot{J}_2 &= F_2 J_\varnothing - (F_1+\gamma_2)J_2 + \gamma_1 J_{1,2}\,,\\ \dot{J}_{1,2} &= F_2 J_1 + F_1 J_2 - (\gamma_1+\gamma_2)J_{1,2}\,,\end{aligned} \tag{S58}$$

where $J_\varnothing = 1 - J_1 - J_2 - J_{1,2}$. If we let $I_1 = J_1 + J_{1,2}$ and $I_2 = J_2 + J_{1,2}$, then model (S58) is equivalent to

$$\begin{aligned}
\dot{I}_1 &= \beta_1(I_1 + \sigma_1 J_{1,2})(1 - I_1) - \gamma_1 I_1\,, \\
\dot{I}_2 &= \beta_2(I_2 + \sigma_2 J_{1,2})(1 - I_2) - \gamma_2 I_2\,, \qquad \text{(S59)} \\
\dot{J}_{1,2} &= \beta_1(I_1 + \sigma_1 J_{1,2})(I_2 - J_{1,2}) + \beta_2(I_2 + \sigma_2 J_{1,2})(I_1 - J_{1,2}) - (\gamma_1 + \gamma_2)J_{1,2}\,.
\end{aligned}$$

Proceeding as in Supplementary Information, Section S1.1, let $P = I_1 I_2$ and $Z = P - J_{1,2}$. Thus,

$$\dot{Z} = -[\beta_1(I_1 + \sigma_1 J_{1,2}) + \beta_2(I_2 + \sigma_2 J_{1,2}) + \gamma_1 + \gamma_2]Z\,. \qquad \text{(S60)}$$

Since the expression inside the brackets is positive, $Z(t) \to 0$ as $t \to \infty$. The prevalence of co-infection by interacting pathogens is asymptotically equal to the product of their prevalences. Therefore, $Z = 0$ does not imply pathogens do not interact.

### S2 Sources of data and side results of model fitting

#### S2.1 Additional fitting of the NiSP model

Results of fitting the NiSP model to data from four publications for strains of a single pathogen, that may plausibly be assumed epidemiologically-interchangeable (López-Villavicencio et al., 2007; Seabloom et al., 2009b; Chaturvedi et al., 2011; Koepfli et al., 2011) are presented in Fig. 3 of the main text. Results for three further data sets concerning different pathogens of a single host (Andersson et al., 2013; Moutailler et al., 2016; Nickbakhsh et al., 2016) are in Fig. S1.

For convenience the raw data as used in model fitting for these additional datasets are re-tabulated in Table S2. Results of model fitting are summarized in Table S3. Ambiguities needed to be resolved in collating these data from what is reported in the original publications. The data presented in Moutailler et al. (2016) are inconsistent, in as much as it is reported that a total of 267 ticks were tested, but the percentage data in the section “Co-infections and associations between pathogens” of the paper instead indicate 262 is the correct total. We have used the value 262 here. Misreporting of the number of uninfected hosts in reference Seabloom et al. (2009b) has been corrected by reference to the original data (Seabloom et al., 2009a) after personal communication with the authors.

| Pathogens with $n$ distinct types, strains or clones | $n$ | Observed counts, $O_k$ | | | | | | | | | | Total |
|---|---|---|---|---|---|---|---|---|---|---|---|---|
| | | 0 | 1 | 2 | 3 | 4 | 5 | 6 | 7 | 8 | 9 | $N$ |
| Pathogens of *Ixodes ricinus* ticks | 37 | 147 | 66 | 24 | 18 | 5 | 2 | - | - | - | - | 262 |
| Barley and cereal yellow dwarf viruses | 5 | 1570 | 224 | 69 | 17 | 6 | 4 | - | - | - | - | 1890 |
| Respiratory viruses | 11 | 17630 | 8568 | 964 | 105 | 15 | 2 | - | - | - | - | 27284 |

Table S2: Sources of data for fitting the NiSP model in which pathogen species, clones or strains are assumed to be epidemiologically-interchangeable. The data sets include pathogens of *I. ricinus* ticks (Moutailler et al., 2016), barley yellow dwarf viruses (Seabloom et al., 2009b), respiratory viruses (Nickbakhsh et al., 2016).

| | NiSP | | Binomial | | | GoF |
|---|---|---|---|---|---|---|
| | $R_0$ | $L$ | $p$ | $L$ | ΔAIC=2ΔL | $p$ |
| Pathogens of *Ixodes ricinus* ticks | 1.021 | -314.1 | 0.020 | -329.3 | 30.5 | **0.476** |
| Barley and cereal yellow dwarf viruses | 1.051 | -1180.8 | 0.048 | -1261.9 | 162.2 | 0.000 |
| Respiratory viruses | 1.037 | -22619.0 | 0.036 | -21731.9 | -1774.2 | 0.000 |

Table S3: Fitting the NiSP model. The NiSP model was highly supported over the binomial model ($\Delta AIC \gg 10$) in all cases tested but one (respiratory viruses), where the binomial model is highly supported over the NiSP model. The final column of the table – GoF – corresponds to the goodness-of-fit test of the NiSP model; values $p > 0.05$ correspond to lack of evidence for failure to fit the data, and so the NiSP model is adequate for the data concerning pathogens of *Ixodes ricinus* ticks (Moutailler et al., 2016).

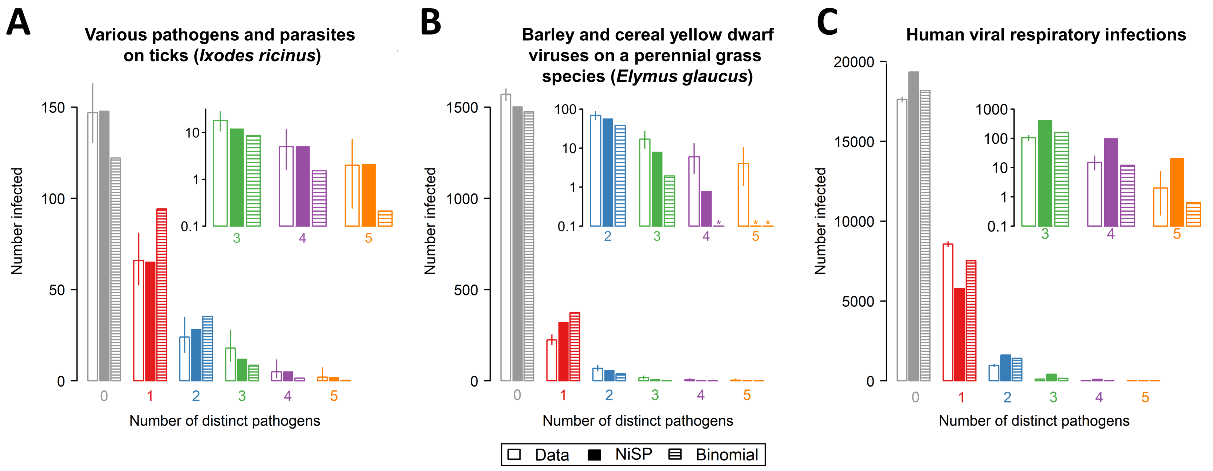


Figure S1: Comparing the best-fitting NiSP model with a binomial model (i.e. statistical independence) for: (A) Pathogens of *Ixodes ricinus* ticks (Moutailler et al., 2016); (B) Barley and cereal yellow dwarf viruses (Seabloom et al., 2009b); (C) Human respiratory viruses (Nickbakhsh et al., 2016). Insets to each panel show a “zoomed-in” section of the graph corresponding to high multiplicities of pathogen co-infection. Asterisks indicate predicted counts smaller than 0.1. For the data shown in (A), there is no evidence that the NiSP model does not fit the data, and so our test indicates the pathogens do not interact. For the data shown in (B), although the NiSP model is a better fit to the data than the binomial model, there is evidence of lack of goodness-of-fit, and so our test indicates these pathogens interact (or are epidemiologically different). For the data shown in (C), although the binomial model is a better fit to the data than the NiSP model, there is evidence of lack of goodness-of-fit, and again it can be concluded that these pathogens interact (or are epidemiologically different).

#### S2.2 Fitting the NiDP model

##### S2.2.1 Sources of data

Howard et al. (2001) report results of analyzing 73 data sets concerning multiple *Plasmodium* spp. causing malaria (rows 68–140 of Table 1 in that paper). We re-analyzed the subset of these studies satisfying the additional constraints that they considered:

- interactions between three *Plasmodium* species (omits 16 rows corresponding to only two pathogens, *viz.* 73, 81, 86, 89-92, 104, 105, 107, 110, 120, 126, 134 and 139-140, as well as 2 rows corresponding to four pathogens, *viz.* 125 and 128);
- disease status of at least 100 individuals (omits 8 rows, *viz.* 72, 85, 87, 115, 129, 135, 136 and 138).

These constraints were imposed simply to reduce the number of studies, rather than because our methodology could not handle such data. We also omitted six of the remaining data sets – rows 83, 93, 94, 121, 122 and 131 – since we found it impossible to unambiguously reconcile the data as reported in the publication to counts of different types of infection. Most often this was because the data were reported as percentages rounded to a small number of significant figures, which did not unambiguously specify the raw number of individuals infected by each combination of pathogens. This left a final total of 41 data sets taken from 35 distinct papers: 24 data sets considering the three-way interaction between *P. falciparum*, *P. malariae* and *P. vivax* (denoted FMV in Howard et al. (2001)) and 17 data sets considering the three-way interaction between *P. falciparum*, *P. malariae* and *P. ovale* (denoted FMO in Howard et al. (2001)). The data sets are re-tabulated for convenience in Table S4.

##### S2.2.2 Recreating the analyses of Howard et al. (2001)

We did not explicitly recreate the analysis based on log-linear models as presented by Howard et al. (2001), since no information was given in the paper on how to handle sampling zeros (i.e. cases in which within an individual data set the count of individual infected by a particular combinations of pathogens is zero). Given the

| | | | Observed counts, $O_\Gamma$ | | | | | | | |
|---|---|---|---|---|---|---|---|---|---|---|
| | | $N$ | ∅ | F | M | V | FM | FX | MX | FMX |
| 74 | Léger et al. (1923) | 250 | 83 | 111 | 49 | 1 | 6 | 0 | 0 | 0 |
| 75 | Bédier et al. (1924) | 135 | 45 | 58 | 27 | 3 | 2 | 0 | 0 | 0 |
| 76 | Knowles and White (1930) | 809 | 642 | 149 | 12 | 1 | 2 | 2 | 0 | 1 |
| 82 (!) | Dorolle (1927) | 652 | 232 | 258 | 64 | 54 | 32 | 12 | 0 | 0 |
| 84 | Phillips (1923)* | 645 | 409 | 112 | 10 | 109 | 0 | 4 | 1 | 0 |
| 88 | Lalor (1913)* | 207 | 94 | 47 | 21 | 40 | 0 | 3 | 2 | 0 |
| 106 | Wilson (1936) | 3393 | 1784 | 1103 | 87 | 19 | 244 | 63 | 2 | 91 |
| 108 (!) | Borel and Levanan (1927) | 1249 | 885 | 227 | 23 | 92 | 3 | 16 | 3 | 0 |
| 109 (!) | Borel and Levanan (1927) | 1022 | 947 | 19 | 12 | 39 | 0 | 4 | 1 | 0 |
| 111 | Banerjea (1930)* | 1519 | 578 | 225 | 7 | 668 | 0 | 41 | 0 | 0 |
| 112 | Khambata (1913)* | 112 | 72 | 26 | 8 | 4 | 0 | 1 | 1 | 0 |
| 113 | Ramsay (1928)* | 1514 | 1073 | 268 | 9 | 160 | 0 | 3 | 1 | 0 |
| 114 | Bailey (1928)* | 1068 | 547 | 396 | 67 | 35 | 18 | 5 | 0 | 0 |
| 116 | Masterman (1913) | 700 | 238 | 317 | 75 | 55 | 4 | 9 | 2 | 0 |
| 117 | Angus (1919)* | 40168 | 28936 | 2614 | 9 | 8483 | 0 | 126 | 0 | 0 |
| 118 | Gordon et al. (1991) | 268 | 208 | 14 | 6 | 30 | 1 | 6 | 3 | 0 |
| 119 | Lalor (1912)* | 151 | 52 | 38 | 13 | 33 | 4 | 6 | 4 | 1 |
| 123 | Carter (1927)* | 11260 | 9510 | 170 | 568 | 986 | 2 | 4 | 19 | 1 |
| 124 | Schnuffer (1938) | 3266 | 2148 | 593 | 234 | 196 | 42 | 30 | 15 | 8 |
| 127 | Banchongaksorn et al. (1996) | 913 | 487 | 221 | 5 | 179 | 0 | 21 | 0 | 0 |
| 130 | Treadgold (1918) | 540 | 396 | 3 | 1 | 136 | 0 | 4 | 0 | 0 |
| 132 | Collins et al. (1988) | 614 | 407 | 151 | 11 | 19 | 3 | 21 | 2 | 0 |
| 133 | Mizushima et al. (1994) | 506 | 231 | 144 | 0 | 81 | 1 | 39 | 4 | 6 |
| 137 | United Fruit Co. (1925)* | 2742 | 1973 | 435 | 14 | 299 | 0 | 21 | 0 | 0 |
| 68 | Campbell et al. (1987) | 147 | 77 | 56 | 0 | 1 | 11 | 1 | 0 | 1 |
| 69 | Campbell et al. (1987) | 142 | 26 | 68 | 3 | 0 | 40 | 4 | 0 | 1 |
| 70 | Campbell et al. (1987) | 196 | 41 | 112 | 2 | 0 | 25 | 10 | 0 | 6 |
| 71 | May et al. (1997) | 230 | 40 | 123 | 1 | 0 | 32 | 7 | 0 | 27 |
| 77 | Alifrangis et al. (1999) | 126 | 5 | 72 | 2 | 0 | 31 | 13 | 0 | 3 |
| 78 | Hellgren et al. (1994) | 163 | 32 | 105 | 5 | 0 | 20 | 1 | 0 | 0 |
| 79 | Thomson et al. (1994) | 1465 | 770 | 641 | 17 | 2 | 30 | 5 | 0 | 0 |
| 80 | Gbary et al. (1988) | 735 | 444 | 234 | 22 | 7 | 20 | 7 | 1 | 0 |
| 95 | Deloron et al. (1989) | 1465 | 770 | 641 | 17 | 2 | 30 | 5 | 0 | 0 |
| 96 | Deloron et al. (1989) | 245 | 130 | 95 | 0 | 0 | 16 | 4 | 0 | 0 |
| 97 | Deloron et al. (1989) | 253 | 126 | 109 | 4 | 0 | 13 | 0 | 0 | 1 |
| 98 | Deloron et al. (1989) | 225 | 136 | 82 | 0 | 0 | 3 | 4 | 0 | 0 |
| 99 | May et al. (1997) | 159 | 97 | 56 | 0 | 1 | 3 | 2 | 0 | 0 |
| 100 | Trape et al. (1992) | 2465 | 2372 | 85 | 6 | 0 | 1 | 1 | 0 | 0 |
| 101 | Trape et al. (1994) | 8539 | 2208 | 4254 | 133 | 50 | 1435 | 227 | 3 | 229 |
| 102 | Molineaux et al. (1980) | 7026 | 2658 | 3295 | 143 | 36 | 742 | 108 | 6 | 38 |
| 103 | Molineaux et al. (1980) | 6526 | 3474 | 2015 | 183 | 15 | 757 | 42 | 2 | 38 |

Table S4: Data sets as extracted from the source references for studies focusing on interactions between *P. falciparum*, *P. malariae* and either *P. vivax* (i.e. FMV) or *P. ovale* (i.e. FMO). The asterisks indicate that the corresponding data sets were extracted from (Knowles and White, 1930). The number in the left-most column shows the number of the relevant row in Table 1 of Howard et al. (2001). The rows with (!) correspond to studies for which the total number of individuals sampled as reported by Howard et al. (2001) do not match what we found on interrogating the original paper; in all cases, we used the corrected values as shown in the table. Note that many of the references are to Knowles and White (1930); this corresponds to cases for which the data from the originally-listed source were extracted from the large compendium collated in 1930 by Knowles & White. The notation X (in FX, MX, or FMX) corresponds either to V (i.e. *P. vivax*, upper part of the table, data sets 74–137) or to O (i.e. *P. ovale*, lower part of the table, data sets 68–103).

small size of many of the studies, this was quite common, affecting 34 of these 41 data sets. The statistical difficulty is that at least one of the models involved in the model selection procedure cannot then reliably be estimated, since an estimated coefficient in a Poisson regression model tends to negative infinity. In turn this means that model selection based on log-likelihood ratio tests breaks down (Fienberg and Rinaldo, 2012). How such sampling zeros affect log-linear models with sparse data sets is an active area of current research in the methodological statistical literature, e.g. (Fienberg and Rinaldo, 2012). It is unclear from what is presented in Howard et al. (2001) precisely how such cases were handled; correspondence with those authors we could contact also did not reveal what precisely had been done in the original analysis (personal communication Prof. Christl Donnelly). We note that, since our methods are based on multinomial sampling rather than Poisson counts, statistical difficulties surrounding sampling zeros simply do not affect our analyses.

#### S2.3 Fitting the models with specific clearance

##### S2.3.1 Fitting the models

All models were fitted after transformation to allow only biologically-meaningful values of parameters, by estimating $\log(\hat{\gamma}_i)$ to ensure only positive values of $\hat{\gamma}_i$ are permissible, and using these to estimate the infection rates after transformation, with $\log\left(\hat{\beta}_i/(\hat{\gamma}_i+1)-1\right)$ (which ensures $R_{0,i}>1$).

However, we noticed that this estimation method may return extremely high values of $\hat{\gamma}$ and $\hat{\beta}$ in the NiSP model. This is because we scaled $\beta$ and $\gamma$ relative to $\mu$, while the optimal value of $\mu$ may be zero. The specific case $\mu = 0$ corresponds to statistical independence (see Sections S1.4.3 and S1.4.5). When the data look statistically independent, the estimation algorithm may diverge. For this reason, we did not explore parameter estimation in the NiDP model with $\gamma > 0$ for the malaria data, as its treatment would have required specific considerations that felt beyond the scope of this paper, since our purpose is not to draw conclusions about interaction among malaria species.

##### S2.3.2 Results of fitting the two-parameter NiSP model

Results of model fitting are summarized in Table S5.

| | NiSP ($\beta$ only) | | NiSP ($\beta$ & $\gamma$) | | | Model selection | | Binomial | | | GoF |
|---|---|---|---|---|---|---|---|---|---|---|---|
| | $\beta$ | $L$ | $\beta$ | $\gamma$ | $L$ | $\chi^2$ | $p$ | $p$ | $L$ | ΔAIC | $p$ |
| Human papillomavirus | 1.032 | -6580.9 | **1.178** | **0.142** | -6573.1 | 7.794 | **0.005** | 0.031 | -6868.8 | 589.3 | **0.986** |
| Pathogens of *I. ricinus* ticks | **1.021** | -314.1 | 1.161 | 0.137 | -313.7 | 0.360 | 0.549 | 0.020 | -329.3 | 30.5 | **0.476** |
| Anther smut (*M. violaceum*) | **1.009** | -611.4 | 1.009 | 0.000 | -611.4 | 0.000 | 1.000 | 0.009 | -690.8 | 158.8 | 0.000 |
| Barley yellow dwarf viruses | **1.051** | -1180.8 | 1.051 | 0.000 | -1180.8 | 0.000 | 1.000 | 0.048 | -1261.9 | 162.2 | 0.000 |
| *Borrelia afzelii* on bank voles | **1.044** | -652.1 | 1.044 | 0.000 | -652.1 | 0.000 | 1.000 | 0.040 | -799.0 | 293.8 | 0.000 |
| Malaria (*Plasmodium vivax*) | 1.021 | -3169.2 | **1.162** | **0.138** | -3164.6 | 4.588 | **0.032** | 0.021 | -3467.3 | 603.5 | 0.000 |
| Respiratory viruses | 1.037 | -22619.0 | $4.7 \times 10^9$ | $4.6 \times 10^9$ | -21731.9 | 887.057 | 0.000 | **0.036** | -21731.9 | -2.0 | 0.000 |

Table S5: Fitting the NiSP model to data sets corresponding to human papillomavirus (Chaturvedi et al., 2011), pathogens of *I. ricinus* ticks (Moutailler et al., 2016), anther smut (*M. violaceum*) (López-Villavicencio et al., 2007), barley yellow dwarf viruses (Seabloom et al., 2009b), *Borrelia afzelii* on bank voles (Andersson et al., 2013), malaria (*Plasmodium vivax*) (Koepfli et al., 2011), respiratory viruses (Nickbakhsh et al., 2016). Parameters for the best-fitting variant of the NiSP model for each pathogen species, strain or clone are highlighted in bold; the two-parameter model is supported in cases for which $p < 0.05$ in the Model Selection part of the table (including human papillomavirus and malaria (*Plasmodium vivax*)). The NiSP model was highly supported over the binomial model (ΔAIC $\gg$ 10) in all cases tested but one (Respiratory viruses). The final column of the table – GoF – corresponds to the goodness-of-fit test of the best-fitting model; values $p > 0.05$ correspond to lack of evidence for failure to fit the data, and so the NiSP model is adequate for the data concerning human papillomavirus and pathogens of *Ixodes ricinus* ticks. These results are qualitatively identical to those for the model without specific-clearance as presented in the main text. Note that in the NiSP model, $\beta$ and $\gamma$ are scaled relative to $\mu$. This is why $\beta$ and $\gamma$ of NiDP reach extremely high values for respiratory viruses. Parameter estimation tends to $\mu = 0$, which actually corresponds to the binomial model, which has one fewer parameter (see Section S1.4.5 and Fig. S2). Hence ΔAIC $= -2$ for Respiratory viruses, since the NiSP model requires one additional parameter compared to the binomial model.

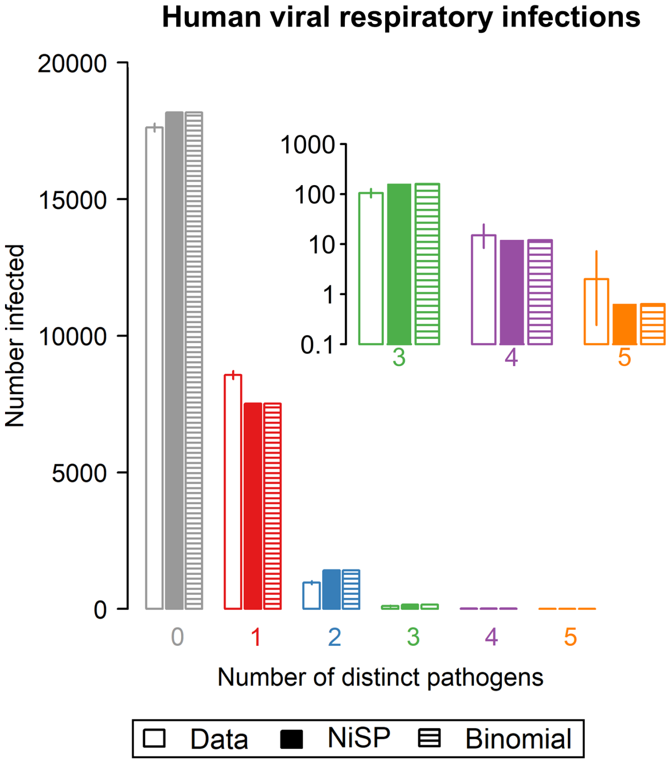


Figure S2: Comparing the best-fitting two-parameter NiSP model with a binomial model (i.e. statistical independence) for human respiratory viruses (Nickbakhsh et al., 2016). Insets to each panel show a "zoomed-in" section of the graph corresponding to high multiplicities of pathogen co-infection. This figure shows that the best-fitting NiSP model converges to the binomial model in this case (which is a special case of NiSP for $\mu = 0$, see section S1.4.5).

Models were fitted by maximum likelihood, with model selection done via $\chi^2$ tests on the likelihood-ratio (Bolker, 2008) or the Akaike Information Criterion (Sakamoto et al., 1986), depending on whether or not models were nested.

##### S2.3.3 Fitting the NiDP model with specific clearance

Estimated parameters occasionally diverge in the more complex version of the NiSP model with specific clearance, and very large numeric values of best-fitting epidemiological parameters can be obtained (but a reasonable value of $R_0$). Exploratory investigations suggested that fitting the NiDP model with specific clearance to the malaria data was affected by this type of identifiability issue, and so would therefore have required a specific treatment. Since our purpose here was not to draw conclusions about interactions among malaria species, but instead to show the utility of our overall approach, we did not pursue this analysis further.
